# Tn7 family bacterial transposons known for tightly controlled target site selection are distributed across Asgard and archaeal groups

**DOI:** 10.1101/2025.07.27.667033

**Authors:** Jordan Thesier, Shan-Chi Hsieh, Richard D. Schargel, Robert J. Wingo, Joseph E. Peters

## Abstract

Tn7 family elements are DNA transposons in bacteria that show tight control of targeting, including multiple varieties that evolved guide RNA-directed transposition. We find diverse Tn7 elements across all archaeal superphyla, including many examples in the Asgard group where the branchpoint with eukaryotes is believed to reside. Mirroring what is known from Tn7 elements in bacteria, specialized specific sites were targeted like tRNA genes and housekeeping genes like ribonucleotide reductase. Reconstituting an Asgard element confirmed that the element possesses the hallmark properties of the Tn7 family. Tn7 elements in bacteria evolve new targeting pathways by recursive cooption of DNA binding domains a process that was also found with protein domains from the eukaryote-archaea sisterhood in the archaeal elements. This work expands our understanding of mobile DNA diversity in archaea and the role a specialized DNA transposon family plays in gene transfer within and between domains.

## Introduction

The archaeal domain shows substantial gene transfer with bacteria and is a sister group with modern day eukaryotes. Genetic transfer between the three domains raises important questions about the origin of genetic systems now found in each branch of life. Mobile genetic elements can be important vehicles for the spread of genetic systems but the details of how this occurs between the domains is unclear. While it is appreciated that major families of mobile elements that are well-studied in bacteria are present in archaea, the important details of some of the most specialized types of mobile elements are lacking. DNA transposons are known to be common across all domains of life and generally move without extensive control over the places where they insert. However, a specialized group within DNA transposons, the Tn7 family elements, tightly control where they insert and only transpose once they have identified a preferred site of insertion. This level of control is manifest in the observation that Tn7 elements evolved guide RNA-directed transposition on multiple independent occasions by coopting CRISPR-Cas systems^1^. Tn7-like elements primarily evolve new targeting modalities by coopting a broad diversity of DNA binding domains that accommodate a variety of DNA sequences that allow the element to adapt to safe integration sites and pathways that promote distribution of the members^2^.

Four transposon-encoded proteins are broadly associated with Tn7-like elements: TnsA, TnsB, TnsC, and TniQ/TnsD^2^. In prototypic Tn7 and most other examples, the transposase is heteromeric, formed by two proteins TnsA and TnsB. The use of two proteins in Tn7-like elements allows transposition to occur by a cut-and-paste process where the element is completely excised from the donor DNA^3,4^. The TnsB subunit of the transposase is a sequence specific DNA binding protein that recognizes binding sites at the transposon ends that flank each side of the element. These flanking ends recognized by TnsB delineate the entire structure that moves as a unit during transposition. Additionally, the transposon ends impart information that allows the machinery to control the position of the insertion and the directionality of integration events^5–7^.

Transposition target sites are typically recognized with TniQ domain-containing proteins that are often fused to a DNA-binding domain. Many different DNA-binding domains have been coopted in a modular process that has evolved to recognize safe sites in the chromosome referred to as attachment (*att*) sites. The TniQ domain interacts with the AAA+ TnsC component which forms a ring or rings around the DNA^8–10^, bridging target recognition to recruitment of the transposase bound to the ends of the transposon^11,12^. Prototypic Tn7 uses ATPase activity to build a TnsC filament in a stepwise fashion, controlling target complex assembly^8^, but how other Tn7 family members use ATPase activity to control transposition appears to vary.

Expanded metagenomic data and advanced techniques for sequence assembly have allowed the diversity of archaea to be more fully appreciated. Archaeal nomenclature is currently unsettled, however, the superphyla that make up archaea are frequently discussed under a growing list of phyla^13^(**Fig. 1**). Much of what we know about archaea comes from unculturable representatives; examples that are known only by DNA sequence are given candidate (*Candidatus - Ca.)* species names. One of the most exciting recent advances in our understanding of archaea is the proposed origin of eukaryotes from within the Asgard group / Asgardarchaeota phylum, likely branching from or closely related to the Heimdallarchaeia lineage^14–16^. Our understanding of mobile elements within archaea is also in its infancy, typically relying on bioinformatic approaches. These involve looking for DNA repeats that are common in these elements, examining CRISPR systems for spacers that match the putative elements^17^, or drawing comparison to known systems in bacteria. However, targeted analyses are still needed to understand the true diversity of mobile elements that facilitate gene flow within the archaeal domain and between the other domains of life.

**Fig. 1.**
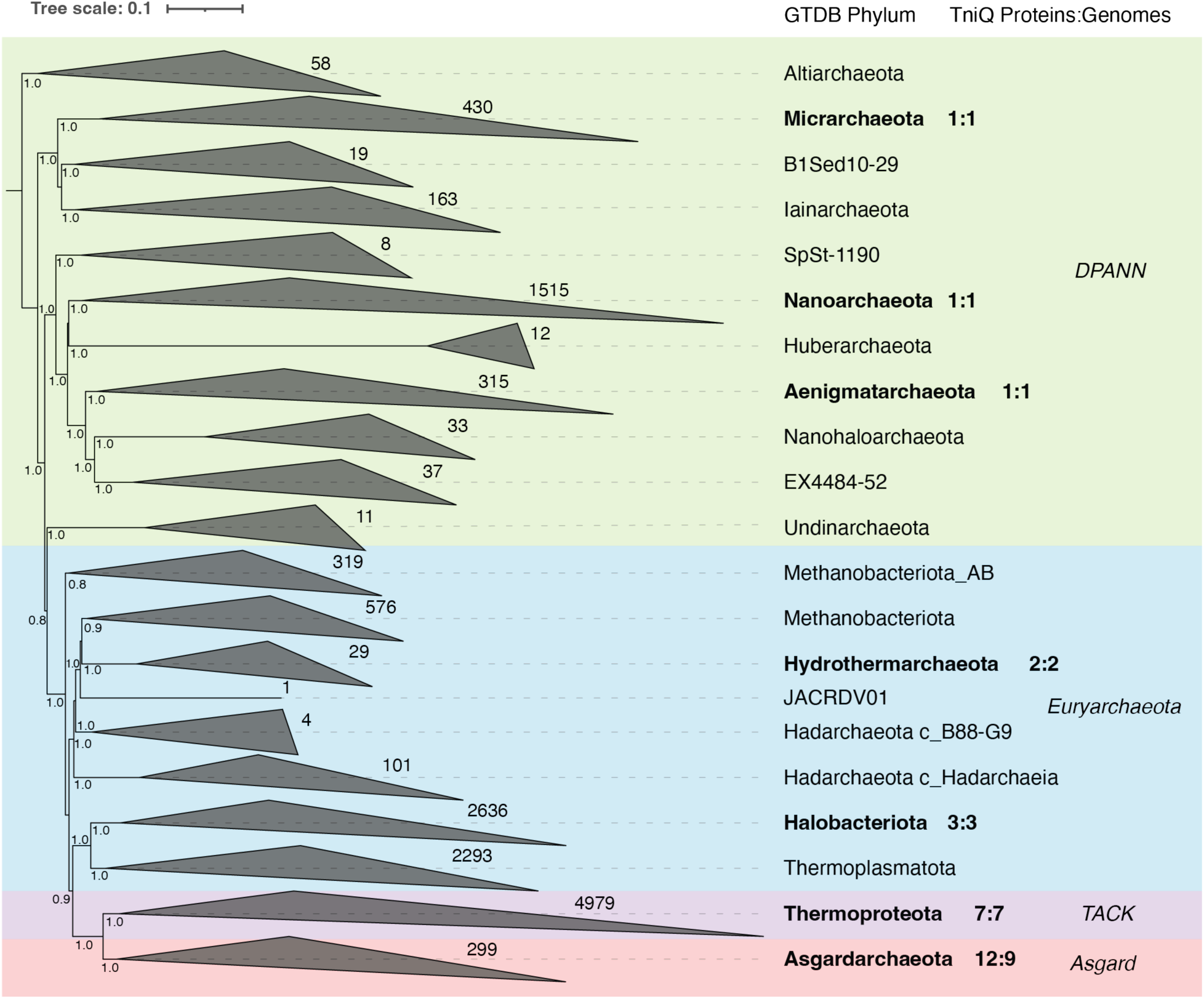
Distribution of TniQ proteins across archaeal assemblies. Similarity tree of archaeal assemblies based on a concatenated alignment of 53 phylogenetic marker genes and taxonomic classification based on the Genome Taxonomy Database (GTDB). Branches are also grouped with under major NCBI superphyla (DPANN, Euryarchaeota, TACK, and Asgard). The tree was artificially rooted with Altiarchaeota. Numbers near the end of collapsed clades represent the number of branches in the clade. Ratios following phylum names represent the number of TniQ proteins identified in relation to the number of TniQ-containing assemblies per phylum. Branch support was calculated with a Shimodaira-Hasegawa test, and values >0.8 are reported at nodes.

In the current work, we identified 10 distinct families of Tn7-like elements that have spread between bacteria and archaea. These families were found in seven different phyla across diverse archaea with several members occurring in Asgardarchaeota, even though these represent a small portion of the total archaeal genome sequences. We found evidence for the same modular adaptation for new attachment sites found in bacteria and in many cases the same types of sites were used, such as tRNA genes and conserved housekeeping genes, like ribonucleotide reductase. In the case of archaea, adaptation occurred via DNA binding domains specific to the archaeal domain with the closest discoverable homologs occurring in metazoans. We reconstituted a Tn7 family element from Asgardarchaeota in a heterologous bacterial host to confirm the functionality and characteristic behaviors of the group. This work expands our understanding of mobile DNA diversity in archaea and the role they play in gene transfer within and between domains.

## Results

### Identification of Tn7-like transposons in archaea

Diverse Tn7-like elements are widespread in bacteria with a growing list of representatives characterized at the molecular level^18–20^. Transfer of genetic information between bacteria and archaea is common, including the exchange of different families of mobile genetic elements, but the distribution of Tn7-like elements in archaea has not been directly examined. To investigate if Tn7 family elements have adapted to archaea, we downloaded 14,044 archaeal genome assemblies from the National Center for Biotechnology Information (NCBI). Using GTDB-Tk^21–32^ for taxonomic classification, we retained only those genomes classified within the domain Archaea, resulting in 13,858 assemblies. We searched these genomes for the Tn7 family signature protein TniQ using the Pfam profile PF06527 and identified 27 TniQ domain-containing proteins across 24 assemblies, all of which are metagenome-assembled genomes (MAGs). This is consistent with the overall dataset, where MAGs comprise 86% of the genomes (11,906 out of 13,858). The 24 TniQ-containing assemblies span seven different archaeal Genome Taxonomy Database (GTDB) phyla, out of the 21 phyla represented by all genomes (**Fig. 1**). Notably, 38% (9/24) of the TniQ-containing assemblies belong to the Asgardarchaeota phylum, while Asgardarchaeota account for only 2% of the overall dataset (300/13,858).

Using MAGs to understand the distribution of mobile elements comes with special risks from possible contamination of the assemblies. As a first step to help guard against misinterpretation, we subjected the 24 archaeal assemblies to quality assessments. One assembly (GCA_017399555.1) was annotated as contaminated in NCBI and removed from further analysis. Another contig (JAWAQQ010000048.1 in GCA_038893775.1) showed an atypical GC content and noted as a possible contaminated assembly by MAGPurify^33^. This assembly was retained based on the idea that transposons are often horizontally transferred and may differ in nucleotide composition from their host genome. As an additional safeguard to support the archaeal origin of these Tn7-like elements, we used MAGPurify to search for single-copy archaeal phylogenetic marker genes^34^. Even though most contigs were small, many also encoded archaeal marker genes and homologs of functional importance (**Table S1**). Specifically, four *tniQ* genes were found on contigs with single-copy archaeal marker genes and seven additional contigs contained genes with high similarity to archaeal homologs (validated by NCBI BLAST).

To estimate MAG completeness and contamination, we used the CheckM2^35^ pipeline. Completeness ranged from 51% to >99% across the assemblies, while contamination levels varied between <1% and 12% (**Table S2**). Although near-complete MAGs are typically defined as ≥90% complete with ≤5% contamination, and medium-quality as ≥70% complete with ≤10% contamination^36^, our focus was limited to the presence of specific transposon genes rather than the overall genomic potential of the host organism. Therefore, we prioritized contamination levels over completeness.

Several assemblies exceeded 10% contamination, a general threshold for potentially problematic MAGs. In cases where *tniQ* genes were supported by the presence of archaeal marker genes, we retained them. Others, particularly orphan *tniQ* genes without accompanying Tn7-like components, were not pursued in further analyses. The final dataset included 23 assemblies with 25 non-redundant (>90% amino acid identity) TniQ proteins (**Table S2**).

### Tn7 family elements transferred between bacteria and archaea on at least ten occasions

Prototypic Tn7 and related elements typically encode multiple protein families, TnsA, TnsB, TnsC, and TnsD/TniQ related gene products, that allow tightly controlled transposition with precise target site selection. Manual inspection of contigs containing TniQ domain proteins revealed 16 examples likely representing archaeal Tn7-like elements, based on the co-occurrence of multiple Tn7 family proteins (**Fig. 2**). Given the extensive diversification among the Tn7 family of elements in bacteria, we constructed similarity trees of TniQ and TnsC proteins to assess the diversity of these archaeal candidates and their relationship to known bacterial counterparts. Each protein was analyzed in two datasets: a smaller set comprised of the archaeal Tn7-like proteins we identified plus 14 established bacterial representatives, and a larger set combining the archaeal proteins with a broad collection of Tn7-like proteins from a previous study^20^.

**Fig. 2.**
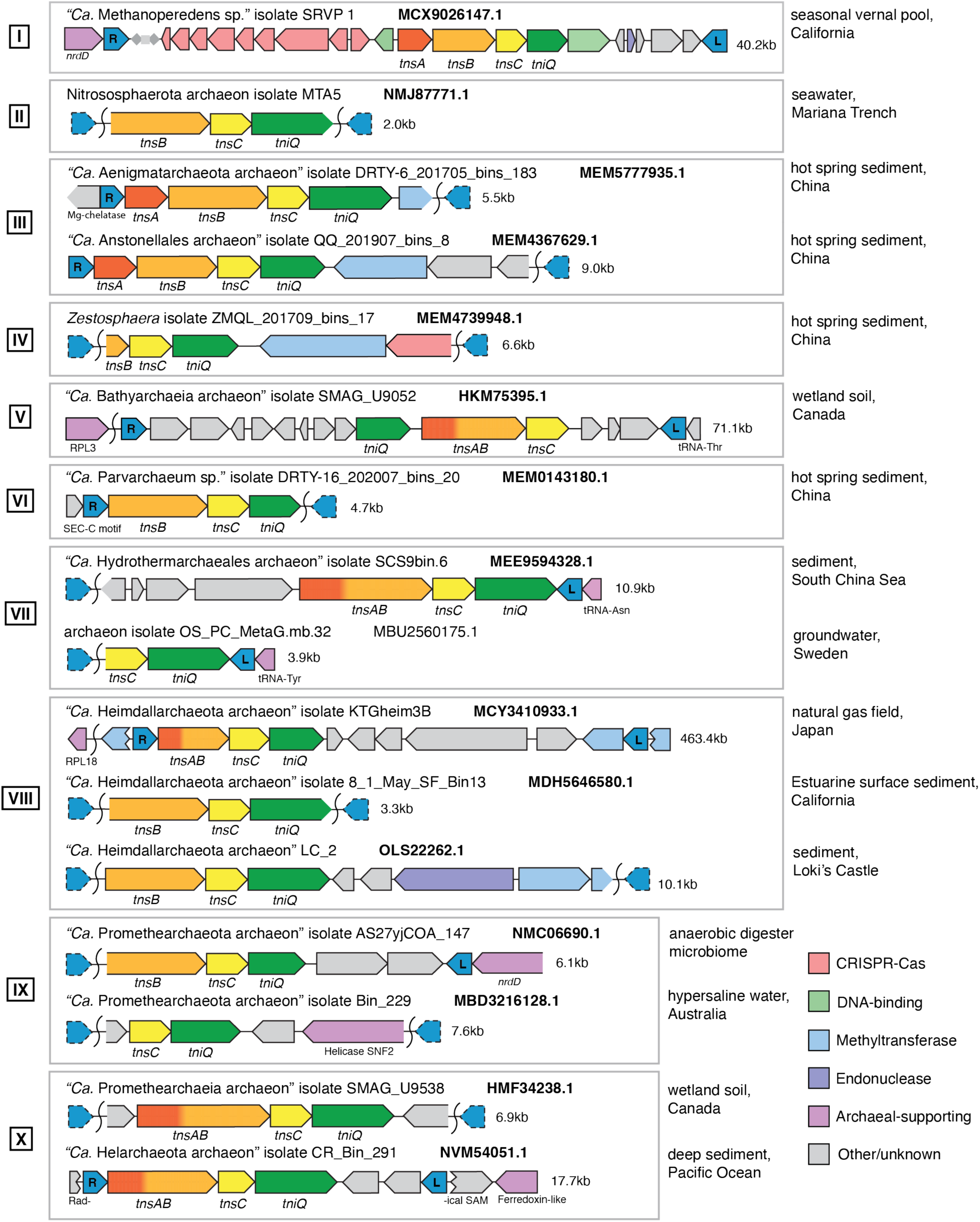
Sixteen candidate archaeal Tn7 family elements. Gene configurations of Tn7-like elements identified on archaeal contigs. Core transposition machinery, predicted cargo, and archaeal-supporting genes are color-coded. Genes truncated by contig boundaries are shown with incomplete outlines. Transposon ends are labeled “R” (right) and “L” (left); dashed outlines indicate undetected ends. Each element is labeled with the NCBI taxonomy of the host archaeon and the TniQ accession number. Contig lengths are shown to the right, and archaeal isolation sources are listed. Roman numerals for clade numbers are noted.

Across both TniQ and TnsC trees (small and large sets), we identified that the 16-candidate archaeal Tn7 family elements grouped into ten distinct clades, each comprising one to three members (**Figs. 3, S1, S2**) In the smaller datasets, archaeal clades were consistently and strongly supported for both TnsC and TniQ (represented by dots on branches). In the larger datasets, archaeal TnsC sequences remained confidently grouped with bacterial examples in all clades except clade VII (**Fig. S1**). The large-set TniQ tree showed less consistent support for archaeal placements, though several clades retained high-confidence with bacterial counterparts, and the same archaeal groupings appeared (**Fig. S2**). Moreover, archaeal sequences within the same clade originated from assemblies classified within a single phylum. One apparent exception involves elements classified as Micrarchaeota and Nanoarchaeota; however, both belong to the same DPANN superphylum, suggesting a possible shared ancestry. Taken together, these findings support a model of 10 independent transfers from bacteria to archaea, followed by lineage-specific diversification in new archaeal hosts.

**Fig. 3.**
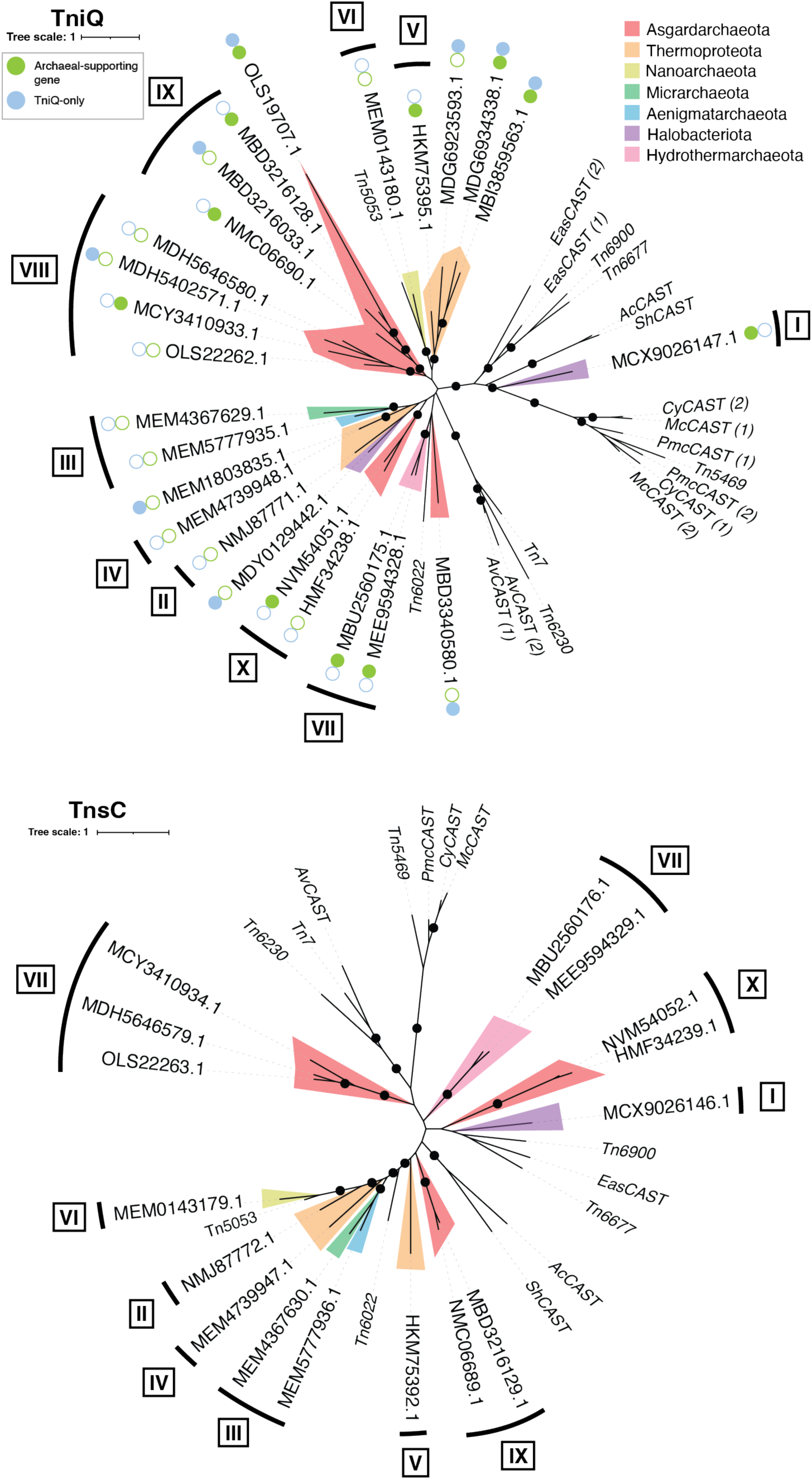
Bioinformatic analysis reveals distinct archaeal clades of Tn7-like proteins. Similarity trees of archaeal TniQ (top) and TnsC (bottom) proteins, each supplemented with 14 characterized bacterial Tn7-like elements. Branches are colored by GTDB phylum. Black dots indicate high support (UFBoot ≥ 70%, SH-aLRT ≥ 0.8, and aBayes ≥ 80%). Roman numerals indicate clade number. For elements encoding multiple TniQ proteins, “(1)” and “(2)” are used to distinguish them. Blue circles mark orphaned TniQ proteins and green circles indicate that the TniQ-containing contig also encodes an archaeal-supporting gene.

In four of the examples (from Clades I, V, VIII and X), we could identify candidate left and right transposon ends suggesting element sizes of 25.8,15.8, 13.9, and 8.4 kb, respectively (**Fig. 2**). The transposition genes encoded in these elements also occurred with the same synteny found with bacterial Tn7 elements. As in bacteria, Tn7 varieties encoding, TnsA and TnsB, with TnsA and TnsB occurring as a single polypeptide, and TnsB without TnsA were identified (**Fig. 2**). The archaeal elements also carried genetic cargo – genes that would move with the element during transposition but were not predicted to be needed for the process itself. Furthermore, we identified nearby host genes positioned at a distance from one end of the element consistent with known bacterial *att* sites, suggesting potential targeting by the transposon (see below).

### Archaeal Tn7 family transposons mirror features identified in bacterial Tn7 family elements

One of the four putative complete elements was identified in “*Ca*. Methanoperedens” (GTDB phylum Halobacteriota, Clade I), in DNAs recovered from seasonal vernal pool sediment in California, USA^37^ (**Fig. 2**). Close relatives of its TniQ and TnsC in similarity trees include type I-F3 CASTs (**Fig 3**)^38^. Notably, this element encodes a type I-D CRISPR-Cas system with components predicted to be a functional defense system^39^ (**Table 1**). While CRISPR-Cas systems have been coopted for guide RNA-directed transposition by Tn7-like elements in bacteria^20^, we find no clear evidence that the element is capable of this targeting mechanism. Canonical CRISPR-Cas defense systems are known to be common in archaea^40^, CRISPR-Cas systems used for guide RNA-directed transposition typically lose the ability to degrade DNA targets and the ability to acquire new spacers on their own^1^, but this element seems to retain these functions through *cas1*, *cas2*, and *csc3*.

**Table 1.** Cargo of putative archaeal Tn7-like elements.

| Clade | Organism | Cargo |
| --- | --- | --- |
| I | "Ca. Methanoperedens sp." isolate SRVP 1 | Type I-D CRISPR-Cas system - cas2 |
|  |  | Type I-D CRISPR-Cas system - cas1 |
|  |  | Type I-D CRISPR-Cas system - cas4 |
|  |  | Type I-D CRISPR-Cas system - cas3 |
|  |  | Type I-D CRISPR-Cas system - cas5/csc1 |
|  |  | Type I-D CRISPR-Cas system - cas7/csc2 |
|  |  | Type I-D CRISPR-Cas system - cas10/csc3 |
|  |  | Type I-D CRISPR-Cas system - cas6 |
|  |  | Type I-D CRISPR-Cas system - csa3 |
|  |  | DNA-binding protein (ArsR superfamily) |
|  |  | DNA-binding HTH domain protein (HTH_17) |
| III | "Ca. Aenigmatarchaeota archaeon" isolate DRTY-6_201705_bins_183 | Methyltransferase (AdoMet_MTases superfamily) |
|  | "Ca. Anstonellales archaeon" isolate QQ_201907_bins_8 | Type I restriction-modification system - DNA methylase subunit (HsdM superfamily) |
| IV | Zestosphaera sp. isolate ZMQL_201709_bins_17 | Type I restriction-modification system - DNA methylase subunit (HsdM superfamily) |
|  |  | CRISPR-Cas system - cas4 |
| VII | "Ca. Hydrothermarchaeales archaeon" isolate SCS9bin.6 | Glycosyltransferase (BcsA) |
|  |  | Ribonuclease (RNase_Zc3h12a superfamily) |
| VIII | "Ca. Heimdallarchaeota archaeon" isolate KTGheim3B | DNA modification methylase (YhdJ) |
|  | "Ca. Heimdallarchaeota archaeon" LC_2 | Type I DNA methyltransferases (HsdM_N superfamily) |
|  |  | Type I restriction-modification DNA methylase (N6_Mtase superfamily) |
|  |  | Type I restriction-modification system - restriction subunit |
| IX | "Ca. Promethearchaeota archaeon" isolate Bin_229 | SNF2 helicase |
|  |  | Beta clamp domain protein |

The Clade I element appears to use an anaerobic ribonucleotide reductase gene (*nrdD*) as an attachment site. A similar arrangement is observed in Clade IX, where a Tn7-like element in the “*Ca*. Promethearchaeota archaeon” is also adjacent to a *nrdD* gene. While the significance of this is uncertain, similar targeting of ribonucleotide reductase genes appears to have evolved independently on multiple occasions. In bacteria, an aerobic ribonucleotide reductase gene, *nrdB*, is the *att* site for a major branch of Tn7 family elements in *Acinetobacter baumannii*^41^. We also identified a group of bacterial Tn7 family elements that independently evolved to target *nrdD,* including *Agathobacter rectalis* AF39-14AC (GCA_003474555.1) and *Fusibacillus kribbianus* YH-rum2234 (GCA_030053595.1) (**Fig. S3**).

In the Clade I element, the TniQ protein may be encoded as a separate polypeptide from the DNA-binding domain responsible for recognizing the *att* site. The TniQ itself is relatively small (<400 amino acids), with a short C-terminal extension beyond the TniQ domain (**Fig. S4**). Directly downstream, however, is a gene encoding a predicted DNA-binding helix-turn-helix protein (HTH) of the HTH_17 superfamily. The closest bacterial homolog of this TniQ protein is also associated with a homologous HTH domain, but in that case, the domains are fused as a single polypeptide in a *Syntrophomonadaceae* host (GCA_012729835.1) (**Fig. S5**). A similar fusion is also found in a different “*Ca*.

Methanoperedens” assembly (GCA_036567565.1), although in this case the gene is cut off by the end of the contig, which likely prevented its detection in our TniQ search. It is possible that the Clade I archaeal sequence erroneously indicates the genes as separate, when in the actual organism they are a single open reading frame; however, there is also precedent for having separate proteins for the TniQ and the DNA-binding components such as in the Tn7 family element Tn6022^41,42^.

The second of the four putative complete elements was identified in a “*Ca*. Bathyarchaeia” assembly (GTDB phylum Thermoproteota, Clade V), recovered from DNA extracted from Canadian wetland soil^43^ (**Fig. 2**). The 15.8 kb element contains fused *tnsAB* genes along with *tnsC* and *tniQ*. It was found adjacent to a threonine tRNA gene that it likely recognizes as an *att* site, given the spacing gene and the end of the element (**Fig. S6**). Similarly, two additional elements (Clade VII) identified in GTBD phylum Hydrothermarchaeota, recovered from South China Sea marine sediment^44^ and Swedish groundwater (BioProject PRJNA627556), also appear to use an archaeal tRNA gene as an *att* site (**Figs. 2, S6**). Targeting of tRNA genes is a recurring feature of bacterial Tn7-like elements as well, including I-D, I-B2, and V-K CAST systems^45–48^.

The third of the four putative complete elements was identified in a “*Ca*. Heimdallarchaeota archaeon” (GTDB phylum Aasgardarchaeota, Clade VIII) from a Japanese natural gas site (BioProject PRJNA491782) (**Fig. 2**). This 13.9 kb element was found on a large contig (>100kb) disrupting an N-4/N-6 DNA methylase gene that was confirmed to be its *att* site (see below). In addition to transposon genes, this element also encodes a methylase gene with 75% amino acid identity to the one it disrupts (**Fig. 4A**). This raises the intriguing possibility that the activity of the disrupted methylase gene is important, and that the element promotes its own persistence by providing a functional replacement. The same contig also harbors genes predicted to encode members of the Rab subfamily of small GTPases, a type of eukaryotic signature protein^49^. Two additional Clade VIII elements were detected from Californian estuary surface water (BioProject PRJNA865744) and sediment around Loki’s castle hydrothermal vents^50^.

**Fig. 4.**
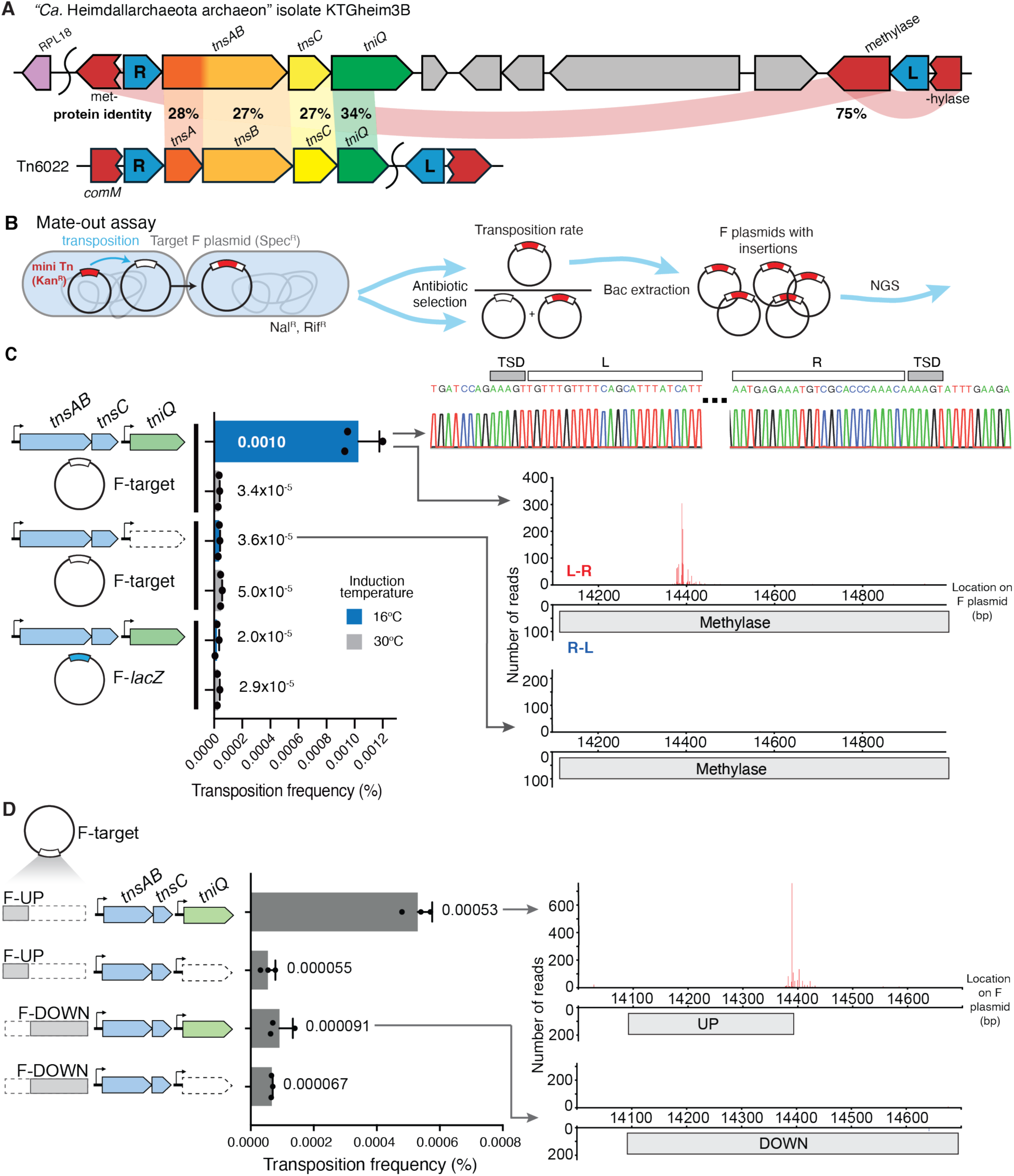
A Tn7-like transposon from “*Ca.* Heimdallarchaeota” is capable of transposition in *E. coli*. **(A)** Gene organization of the putative ”*Ca.* Heimdallarchaeota” transposon compared to the known Tn7-like element Tn6022. Transposition genes are labeled, with transposon ends marked as “R” (right) and “L” (left). Protein identities are marked in color-coded connections. **(B)** Schematic of the mate-out assay used to determine transposition frequency (plasmids expressing transposon genes are omitted for clarity). **(C)** *Left*: Transposition frequencies of the archaeal element into an F plasmid carrying either the target methylase gene (F-target) or *lacZ* (F-*lacZ*). Trials without *tniQ* are indicated as white genes with dashed outlines. Proteins were induced at 30 or 16 °C with 0% arabinose and 0.1mM IPTG. Data are shown as mean + SD, n = 3. *Right, top*: Sanger sequencing of an on-target insertion with TnsABC and TniQ. Target site duplications (TSDs) and transposon ends (L: left end; R: right end) are indicated. *Right, bottom*: Corresponding insertion distributions into F-target via Illumina sequencing., comparing TnsABC + TniQ vs TnsABC alone. Red bars indicate insertions with a L-R orientation, and blue bars indicate insertions with a R-L orientation. **(D)** Testing the archaeal transposon’s target site within the methylase gene. *Left*: The target methylase gene in F-target was truncated to include either the upstream region and TSD (F-UP) or downstream region and TSD (F-DOWN). Transposition frequencies were calculated by mate-out assay. Data are shown as mean + SD, n = 3. *RIGHT*: Corresponding insertion distributions into F-UP and F-DOWN with *tniQ* via Illumina sequencing.

The final putative complete element was identified in a “*Ca*. Helarchaeota” assembly (GTDB phylum Agardarchaeota, Clade X), derived from Pacific Ocean sediment^51^ (**Fig. 2**). This element was also found disrupting a gene, in this case encoding a putative radical SAM associated with replication, recombination, and repair. Homologs of a concatenated version of this protein appear to be archaeal-specific based on NCBI protein BLAST, but the element itself does not encode a known functional homolog. This is also the only archaeal element identified with *tga* transposon ends instead of the more common *tgt* ends, though *tga* ends have been observed in bacterial Tn7-like elements (e.g. Tn7005^52^). The second Clade X element was found in a “*Ca.* Promethearchaeia archaeon” isolated from wetland soil in Canada^43^, and both encode fused *tnsA* and *tnsB* genes along with *tnsC* and *tniQ* genes.

### Adaptation of TniQ to Archaea by cooption of domain-specific DNA-binding domains

TniQ proteins in bacteria are highly modular and are most different in their C-terminal regions associated with DNA-binding and target selection. One of the mechanisms that allow Tn7 family elements to evolve to target new sites is by swapping DNA-binding domains (DBDs). Bacteria, Archaea, and Eukarya are known to have favored repertoires of DBDs^53^, and archaeal biology is often described as a blend of bacterial- and eukaryotic-like features due to their unique evolutionary position between the two domains of life. Archaeal transcription is an example of this hybrid nature, where their basal transcription machinery resembles that of eukaryotes^54–56^, but their specific transcription factors are often more similar to those of bacteria^57–59^. Based on this, we expected the TniQ proteins found in archaea to follow similar trends in their DBDs, being bacterial-, eukaryotic-, or uniquely archaeal-like, if they had indeed adapted to this new domain.

To establish trends with TniQ DBD cooption in the candidate archaeal Tn7 elements, we examined homologs of the archaeal TniQ DBDs using NCBI protein BLAST and report the top 10 hits (**Table S3**). Among the 16 archaeal Tn7 family elements, six TniQs showed homology to eukaryotic proteins, four to both archaeal and bacterial proteins, four to only archaeal proteins, and one to only bacterial proteins. One DBD was too small to analyze. Surprisingly, many of the eukaryotic homologs contained Cys_2_-His_2_ zinc finger domains (C2H2-ZnFs), a DNA-binding motif found in the largest class of eukaryotic transcription factors^60–62^, a configuration that is nearly absent in bacteria^63,64^, and poorly characterized in archaea^65,66^. For example, the DBD of TniQ protein MBD3216128.1 shares 43% amino acid identity to a gastrula ZnF protein (XlCGF8.2DB-like, XP_041843849.1) from *Melanotaenia boesemani*, a species of rainbowfish. These eukaryotic-like TniQ proteins are distributed across Clades VII, VIII, IX, and X, and originate from Hydrothermarchaeota (2 of 6) and Asgardarchaeota (4 of 6) lineages. Furthermore, several of these TniQ DBDs also aligned with their eukaryotic homologs based on AlphaFold3-predicted structures (**Fig. S7**).

To move beyond association and assess whether these proteins truly encode C2H2-ZnFs, we used the Pfam profile PF00096 with subsequent motif validation. We confirmed the presence of C2H2-ZnFs in five archaeal TniQs and detected zero among thousands of bacterial TniQs^42^. Each contained tandem repeats generally following the canonical motif (CX_2-4_CX_12_HX_2-6_H), consistent with the multi-finger arrangement of eukaryotic C2H2-ZnF proteins^63,67,68^ (**Fig S8)**. Although some motifs were degenerate, featuring substitutions in zinc-coordinating residues or partial patterns, such variation is also observed in eukaryotes^69,70^. To further probe the eukaryotic nature of these domains, we queried the archaeal TniQs against HMM profiles of 37 C2H2-ZnF families^71^. One protein, MBD3216128.1, had significant hits with five of these families (MTF1, GFI, FEZ, ZXD, and ZFP362), all of which are associated with the expansion of C2H2-ZnFs in metazoans. Likewise, MBD3216128.1 has the most fingers of the archaeal set and its linkers between fingers are most similar to canonical TGERP, both hallmarks of C2H2-ZnFs in higher eukaryotes^68,71^. The DBD of MBD3216128.1 also aligned with *Homo sapiens* representatives of these five C2H2-ZnF families based on AlphaFold3-predicted structures (**Table S5, Fig. S9**).

Our examination of TniQ DBDs also revealed other signs of potential adaptation. The Clade I element, described above, encodes a TniQ homologous to proteins from both bacterial (*Agathobacter rectalis* AF39-14AC and *Fusibacillus kribbianus* YH-rum2234) and archaeal (another “*Ca*. Methanoperedens archaeon”) sources (**Fig. S5**). Although this element appears to target a *nrdD* gene, its DBD does not resemble those of known bacterial *nrdD*-targeting Tn7 family elements. Clade VII elements also deviate from known bacterial systems: while predicted to target tRNA genes, their DBDs lack homology with bacterial tRNA-targeting systems such as I-D, I-B2, and V-K CASTs, and instead encode C2H2-ZnFs. These differences in DBD structure, despite conserved target types, support the idea that TniQs from bacteria have adapted to different host environments by acquiring new DBDs common to the eukaryote-archaea sisterhood.

Conversely, the presence of highly similar DBDs across archaeal and bacterial elements may not be a sign of adaption, but instead a sign of potential transfer of Tn7-like elements between the domains. The Clade III element from a ”*Ca*. Aenigmatarchaeota archaeon” and the Clade IV element from a *Zestosphaera* sp. encode homologous TniQ proteins that are also similar to a group of bacterial TniQs, including WP_022670411 from *Hippea alviniaea* (**Fig. S10**). Taken together, these results support new DBD cooption by archaeal TniQ proteins, a subset including eukaryotic-like C2H2-ZnFs, and highlight the potential of horizontal gene transfer of Tn7 family elements across domains.

### Reconstitution of targeted transposition by the Tn7-like element from “Ca. Heimdallarchaeota”

To determine if the putative archaeal Tn7-like elements are capable of transposition and if they share known characteristics of Tn7 family elements, we tested the Clade VIII element from “*Ca.* Heimdallarchaeota archaeon” isolate KTGheim3B using a mate-out assay in *E. coli* (**Fig. 4**). In this assay, the transposon genes were optimized for bacterial expression and expressed from plasmids. The DNA sequence of the host methylase gene surrounding the native insertion was cloned into a mobile F plasmid to be tested as a potential *att* site (F-target). A mini transposon comprising an antibiotic resistance gene flanked by the putative transposon ends was situated in an *E. coli* plasmid. Transposition was monitored by inducing expression of the candidate transposon genes and then mating the F plasmids (containing the *att* site) into a recipient strain. Successful transposition was detected by selecting for F plasmids with and without the genetic marker encoded on the mini transposon.

Transposition of the archaeal element was successfully reconstituted in *E. coli* and required the TnsAB, TnsC, TniQ proteins as well as the methylase gene as an *att* site (**Fig. 4C**). Transposition was more frequent at 16°C than 30°C, and at lower protein induction levels (**Fig. S11**). We suspect that lower expression and temperature facilitated folding of the proteins or assembly of nucleoprotein complexes in the non-native host. DNA sequencing confirmed the 5 bp target site duplication (TSD) that is indicative of transposition with Tn7-like transposons (**Fig. 4C**). Using next-generation sequencing (NGS), we observed that insertions frequently occurred at the same exact position and in the same orientation as identified in the archaeal assembly, mirroring the target control found with bacterial Tn7-like elements (**Figs. 4C, S11B**). Specifically, 74.7% of insertions occurred within a 50 bp window surrounding the native TSD in the presence of TniQ, whereas no insertions were detected in this window when TniQ was omitted. Experiments with three other elements were attempted but we were unable to reconstitute transposition in *E. coli* with these examples (**Fig S11**), potentially due to dependence on archaeal-specific host factors that are required in the targeting complex, as protein chaperones, or as DNA bending proteins.

Work with prototypic Tn7 has indicated that the TniQ protein interacts with a sequence adjacent to one side of the attachment site^72^. To confirm this in our archaeal Tn7-like element, we trimmed the target methylase gene located on the target F plasmid to either (a) upstream and including the native TSD (F-UP), or (b) downstream and including the native TSD (F-DOWN). In a mate-out assay, transposition efficiency was higher with F-UP compared to F-DOWN and on-target insertions were only detectable in F-UP via NGS (**Fig. 4D**). Moreover, electrophoretic mobility shift assays (EMSAs) using purified TniQ protein showed a band shift indicative of TniQ-DNA binding only with the upstream methylase gene fragment and not the downstream one (**Fig. S12**). This supports the presence of a specific sequence being recognized in the attachment site in the upstream region of the target methylase gene.

### The archaeal transposon exhibits Tn7 family characteristics

In addition to exhibiting TniQ-mediated targeted transposition, the archaeal Tn7-like element displays other hallmark properties of bacterial Tn7 family transposons, including target site immunity and ATPase-dependent regulation. Target site immunity inhibits a second transposition event from occurring into an *att* site that already contains an insertion^73^. To look for evidence of target site immunity, we cloned an immobile mini transposon in the F plasmid methylase gene at the native TSD (F-target::Tn) to mimic a previous insertion (**Fig. 5A**). This mini transposon contained a different resistance gene and had mutations at the final two base pairs that render it unable to transpose^74^. Consistent with target immunity, TniQ-directed transposition into F-target::Tn was reduced compared to F-target (**Fig. 5A**). Sequencing revealed that most insertions occurred outside the methylase gene, and none were observed at the native insertion site (**Figs. 5B, S11C)**.

**Fig. 5.**
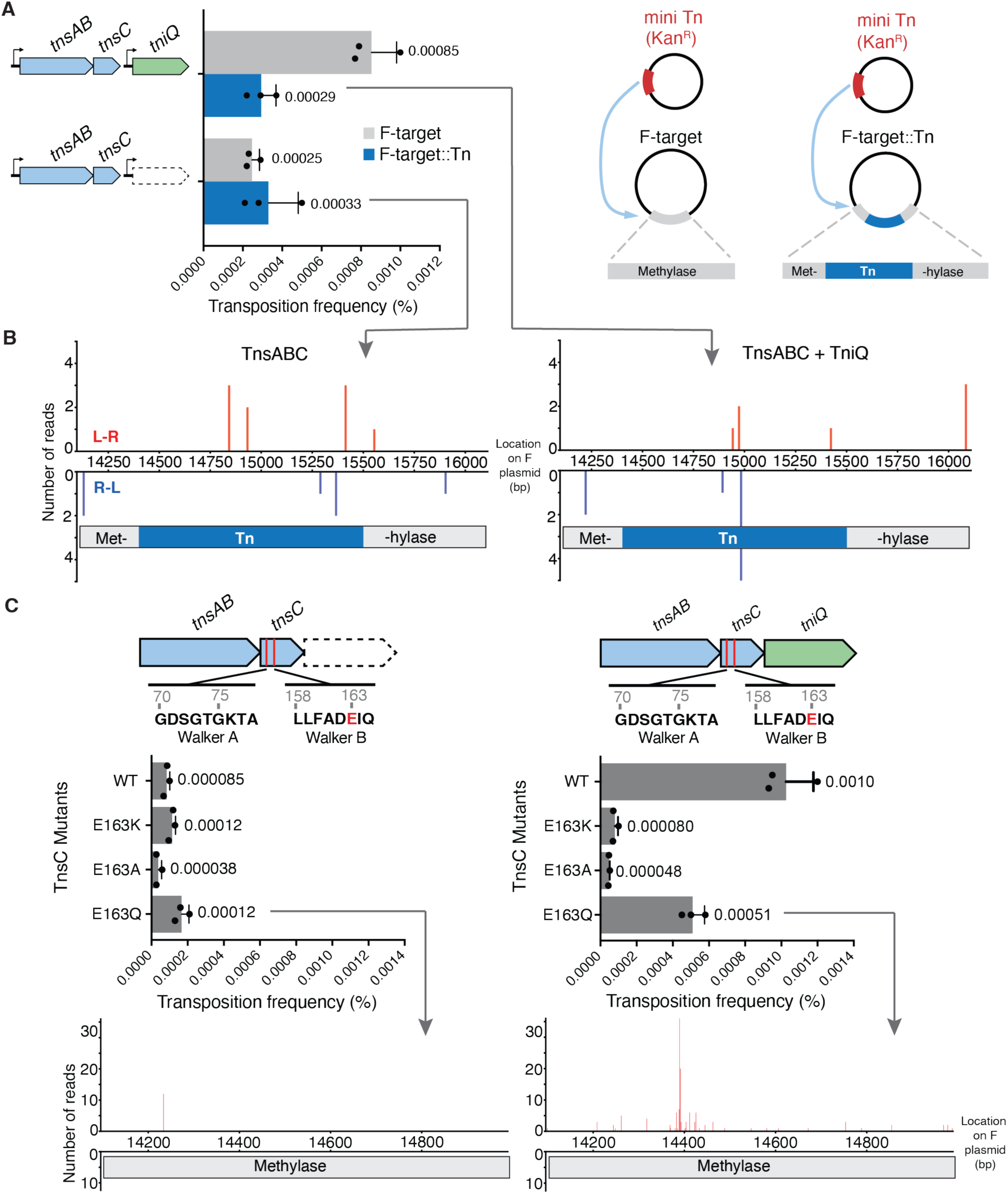
The “*Ca.* Heimdallarchaeota” transposon exhibits hallmark features of Tn7-like elements. **(A)** *Right*: To test for target-site immunity, an immobile version of the archaeal mini element was cloned into the native insertion site within the methylase gene (F-target::Tn). Transposition frequencies into F-target and F-target::Tn were measured using the mate-out assay. *Left*: Transposition frequencies testing target-site immunity. Data are shown as mean + SD, n = 3. **(B)** Corresponding insertion distributions into F-target::Tn based on Illumina sequencing. *Left*: TnsABC-only transposition; *Right*: TnsABC + TniQ transposition. No insertions occur at the native site. **(C)** To assess the role of ATPase activity, the Walker B motif of the archaeal element was mutated. The key glutamate residue responsible for ATP hydrolysis (E163) is indicated in red. Transposition frequency was calculated by mate-out assay, n = 3. Data are shown as mean + SD. Insertion distributions into the target methylase gene are shown below. The E163Q mutant retains limited ability to transpose site-specifically with TniQ.

In addition to target site immunity, we examined the role of ATPase activity on the archaeal Tn7-like element. TnsC is a AAA+ protein that is maintained across Tn7 family elements, but differences are found with how the ATPase activity is used in the diverse members of the family. In prototypic Tn7 and some other cases, mutations in the Walker B motif of Tn7 show a gain-of-function phenotype^9,75–77^, causing transposition to no longer be dependent on TniQ. These insertions occur across DNAs in the cell without a distinct bias. In other cases, mutations in the Walker B motif abolish or substantially reduce transposition^45,78^. We tested different Walker B mutations of the archaeal TnsC in a mate-out assay and found that loss of ATPase activity did not result in a gain-of-function phenotype; instead, these mutations produced negligible transposition levels with or without TniQ (**Fig. 5C**). Mutation of glutamic acid to lysine or alanine in the Walker B motif nearly abolished transposition. Mutating glutamic acid to glutamine, however, retained a low frequency of transposition with a small portion of insertions remaining on-target (**Fig. 5C**). Further experiments will be needed to understand the exact role of ATPase activity in the transposition system, but the genetic results suggest ATPase activity is involved in controlling transposition beyond working as a signaling component during assembly.

## Discussion

In our analysis of archaeal genomes, we identified Tn7 family signature proteins across seven diverse archaeal phyla (**Fig. 1**). We found evidence for at least 10 instances of transfer between bacteria and archaea. This inference is supported by the consistent clustering of archaeal TnsC and TniQ homologs among bacterial relatives in multiple similarity trees, with taxonomically related archaea forming clades within each tree (**Figs. 3, S1, S2**). These patterns suggest that Tn7-like elements were acquired independently by distinct archaeal lineages and subsequently diversified within those groups. Almost half (7 out of 16) of these archaeal Tn7-like systems belong to the Asgardarchaeota phylum, which is specifically intriguing due to the phylum’s close evolutionary relationship to Eukarya^79^.

To better understand how Tn7-like elements may function in archaea, we examined the mechanisms they may use to recognize integration sites. TniQ proteins are highly modular in bacteria, enabling the evolution of targeting diverse *att* sites. We found that some archaeal Tn7-like elements appear to have independently re-evolved targeting to gene classes commonly used by bacterial elements. Specifically, two archaeal elements are predicted to target *nrdD* and three are predicted to target tRNA genes (**Fig. 2**), both of which are *att* sites identified with bacterial Tn7-like elements and other mobile elements such as bacteriophages and Integrating Conjugating Elements. However, the DBDs presumed to mediate this targeting differ substantially between archaeal and bacterial TniQs, supporting the idea that these pathways evolved de novo. None of the archaeal TniQ proteins shares detectable homology with the predicted DBDs of their bacterial counterparts. For instance, two archaeal tRNA-targeting TniQs (Clade VII) contain C2H2-ZnFs, which are entirely absent from bacterial TniQs (**Fig. S8**). Among the archaeal *nrdD*-targeting elements, one encodes a putative accessory DNA-binding protein with no clear homology to bacterial *nrdD*-targeting DBDs, while the other has only a minimal C-terminal extension, suggesting it may lack a DBD entirely (**Fig. S4**). These findings suggest that while archaeal and bacterial Tn7-like elements may converge on similar targets, molecular mechanisms for recognizing those targets are different.

Among these divergent targeting mechanisms, several are especially notable for their prevalence across the archaea-eukaryote sisterhood. We discovered that five archaeal TniQ DBDs encode C2H2-ZnFs, with a majority belonging to Asgardarchaeota (**Fig. S8**). C2H2-ZnFs are one of the largest transcription factor families in eukaryotes and are far more commonly found in eukaryotes than in archaea or bacteria^60–63^. Moreover, the closest homologs to these archaeal DBDs are found in metazoans (**Table S3**).

Since there is growing evidence for eukaryotic origins within the archaeal domain, specifically within the Asgardarchaeota phylum^79^, this shared ancestry raises the possibility that archaeal Tn7-like elements may have evolved features more readily compatible with eukaryotic systems. While Tn7-like elements have not been identified in eukaryotes, the suggestion has been made the Tn7 domains may have been the starting templates for some proteins in eukaryotes^80^. Multiple examples were found where the closest homologs for target recognition DNA binding domains were in humans suggesting they could serve as promising scaffolds for adapting Tn7 family systems as gene editing tools for human therapeutics. On a broader level, the diversity represented in this work should inform the development of genetic tools for a variety of important systems in archaea and eukaryotes.

The strongest support for the activity of these archaeal elements, and their potential for moving across domains, comes from our reconstitution of transposition using an element from ”*Ca.* Heimdallarchaeota” (GTDB phylum Asgardarchaeota, Clade VIII). To our knowledge, this marks the first experimental demonstration of a Tn7-like element derived from the archaeal domain. Remarkably, this element was fully functional in the heterologous bacterial host *E. coli*. We observed site-specific insertion into a favored genomic location in a single orientation, with sequence recognition occurring on one side of the integration point (**Fig. 4C,D**). Additionally, we showed that this archaeal transposon displays target immunity that may use ATPase activity as known from bacterial representatives (**Fig. 5**). These results underscore that core features of bacterial Tn7 family behavior are likely preserved in archaea.

The genomic context of the archaeal element provides further insight into how these systems may be maintained in archaeal hosts in some cases. In most cases in bacteria, TniQ proteins recognize the coding sequence of essential genes, but insertions are directed outside the gene a feature that would allow the resulting integration event to avoid disrupting gene function. We found similar examples of this in archaea with tRNA and *nrdD*-based *att* sites. In contrast, in one case we identified the archaeal element that appears to target and disrupt a methylase gene. However, as it also carries a methylase with 75% amino acid identity as cargo (**Fig. 4A**), we suspect that the native methylase gene is important, but that its function can be replaced by the methylase encoded within the Tn7-like element. In some cases, bacterial Tn7-like elements have also evolved to recognize a position that specifically inactivates the gene they recognize. For example, Tn7-like elements and other integrating elements have evolved to target genes needed for natural transformation^41,42,45^. Presumably, inactivating competence benefits integrating elements by reducing the risk of foreign DNA uptake that could eliminate the element through homologous recombination. By disabling a host gene and replacing its function, the archaeal element may act like an addiction system, ensuring its own persistence. A similar mechanism where a transposon replaces the function it insertionally inactivates has been previously suggested for resolvase activity with Tn5053 and telomere maintenance with telomeric transposons^77,81^. Although both transposon ends could not be identified in many of the archaeal systems, likely due to the relatively short contig sizes, the nearby presumed cargo genes that could be annotated were mostly defense-related, consistent with patterns seen in bacterial Tn7-like elements (**Fig. 2**, **Table 1**).

Altogether, these findings expand our understanding of Tn7-like elements and their functional potential across domains, emphasizing that behaviors characterized in bacterial elements are also likely occurring in archaea. Moreover, the modularity and adaptability observed here highlight the potential of archaeal transposons for genome engineering. It will also be interesting to see how the examples of Tn7 family elements grow as the number of complete archaeal genome assemblies accumulate to provide a better picture of how these elements evolved

## Acknowledgments

We thank Stephen Zinder for his comments on initial work, and the members of the Peters lab, current and former, for helpful discussions.

## Funding

This work was supported by National Institutes of Health grant GM152260 (J.E.P.).

## Author Contributions

Conceptualization: S.-C.H., J.T., J.E.P. Bioinformatics: J.T., S.-C.H., Cloning: R.J.W., S.-C.H., J.T., Experimentation: J.T., S-C.H., R.D.S., J.E.P.

Visualization: J.T., S.-C.H., J.E.P. Writing – original draft: J.T., S.-C.H., J.E.P. Writing – reviewing: J.T., S.-C.H., R.D.S., J.E.P. Supervision and funding acquisition: J.E.P.

## Competing Interests

Cornell University has filed patent applications unrelated to work in this manuscript with some of the authors as inventors.

## Data and materials availability

ll data are available in the main text or supplementary materials.

## Supplementary Materials

### Materials and Methods

#### Bioinformatic discovery of Tn7-like transposons in Archaea

Annotated nucleotide (14,044) and protein (14,042) FASTA files for archaeal genomes (taxid 2157) were downloaded from the National Center for Biotechnology Information (NCBI) GenBank and RefSeq databases via FTP in September 2024. To identify Tn7-like homologs, Hidden Markov Models (HMMs) for TniQ (PF06527) and TnsC (PF13401) were obtained from the European Bioinformatics Institute (EMBL-EBI) Pfam database, and searched against the protein sequences using *hmmsearch* (HMMER v3.4). This yielded 27 TniQ and 17 TnsC proteins. Assemblies were subjected to quality assessment (see below), and 1 assembly with both TniQ and TnsC was excluded due to quality concerns (GCA_017399555.1). To reduce redundancy, proteins with >90% identity were clustered using CD-HIT v4.8.1, resulting in a final set of 25 TniQ and 16 TnsC proteins (**Table S2**).

To assess phylogeny, we supplemented our archaeal dataset with 14 specific Tn7-like elements previously described, or with a large-scale survey of Tn7-like elements by Faure et al., 2023. Protein sequences were trimmed to TniQ or TnsC domains using coordinates from the prior *hmmsearch* –domblout output and aligned with *hmmalign* (HMMER). Maximum-likelihood similarity trees were built with IQ-TREE (v2.2.2.6) using the best-fit substitution model (automatically selected for the large dataset and applied to the smaller one). Branch support was evaluated with 1000 ultrafast bootstrap and SH-aLRT replicates and approximate Bayes tests; branches meeting thresholds of UFBoot ≥ 70%, SH-aLRT ≥ 0.8, and aBayes ≥ 80% were marked on the trees (**Figs. 3, S1, S2**).

#### Assembly Quality Assessment

Taxonomic classifications were assigned to downloaded assemblies using GTDB-Tk *classify_wf* (v2.4) based on the Genome Taxonomy Database (GTDB)^21^. Assemblies classified under the domain Archaea were kept for further analysis (13,858 of 14,044 total). A maximum-likelihood similarity tree was constructed using a concatenated multiple sequence alignment of 53 archaeal single copy marker genes from GTDB-Tk, with FastTree (v2.1.11) and rooted with Altiarchaeota (**Fig. 1**). Nearly all assemblies fell within their expected taxonomic clades, with only 8 branches (0.058%) falling outside (none of which contained a TniQ protein).

To assess the quality of TniQ-containing MAGs, we used MAGPurify (v2.1.2) to identify discordant contigs based on clade- and phylogenetic-specific markers, tetranucleotide frequency, and GC content. Genome completeness and contamination were estimated with CheckM2 (v1.1) (**Table S2**). To support the archaeal origin of TniQ-containing contigs, we searched for “archaeal-supporting” genes defined as single-copy archaeal marker genes identified by MAGPurify and other functionally important genes with high similarity to archaeal homologs. These genes were validated using NCBI’s Basic Local Alignment Search Tool (BLAST) (**Table S1**).

#### Analyzing elements in archaeal assemblies

Gene neighborhood analyses of putative archaeal Tn7-like elements were done on NCBI. Left and right ends were identified based on proximity to Tn7-related genes or known bacterial targets and the presence of TnsB binding motifs detected with MEME Suite^82^. Genes located within 50bp from a left or right end were considered potential targets. Functions of genes were analyzed with NCBI”s Batch Conserved Domain Search and eggNOG-mapper (v2)^83,84^ (**Tables S5, S6**). Protein domains were predicted with InterProScan using the Pfam, PANTHER, and SMART databases (**Figs. S3, S4**).

#### TniQ adaptation analysis

To analyze regions beyond the conserved TniQ domain, archaeal TniQ protein sequences were trimmed to retain everything after this domain. Homologs were identified with protein BLAST and the top 10 hits per protein were reported. Significant homology was considered with E-values < 1e-06. To detect C2H2-ZnF domains, the HMM PF00096 from EMBL-EBI was scanned across archaeal TniQ sequences with *hmmsearch*. Significant hits were defined as those spanning most of the HMM profile (23 residues), with domain E-values < 1e-03, and located adjacent to other hits in the same protein. Motifs following CX_2-4_CX_12_HX_2-6_H were manually checked (**Fig. S8**). Additionally, protein FASTA files for 37 C2H2-ZnF families from Seetharam and Stuart, 2013 were aligned with MUSCLE and used to build HMMs via *hmmbuild* (HMMER). These profiles were used to scan archaeal TniQ proteins, with significant hits defined by full-sequence and domain scores > 100 and E-values < 1e-03. Human representatives of 5 C2H2-ZnF families were used from this study”s dataset for predicted structure alignments (**Fig. S9**) and their *Homo sapiens* origin was confirmed via BLAST (**Table S4**).

#### Bacterial strains and plasmids

Transposition donor strains were constructed in *E. coli* strain BW27783. Plasmids described in supplementary Tables S7 and S8 were synthesized by Twist Bioscience or constructed with manufactured gene fragments by standard molecular cloning techniques and confirmed via DNA sequencing.

#### Growth conditions

*Escherichia coli* strains (**Table S7**) were grown in lysogeny broth (LB), M9 minimal media, or on LB agar supplemented with the following concentrations of antibiotics and supplements where appropriate: 100 μg/ml carbenicillin (Carb), 30 μg/ml chloramphenicol (Cam), 50 μg/ml kanamycin (Kan), 50 μg/ml spectinomycin (Spec), 20 μg/ml nalidixic acid (Nal), 100 μg/ml rifampicin (Rif), 100 μg/ml trimethoprim (Tmp), 0.2% glucose, 0/0.002/0.2% arabinose, 0.2% maltose, and 0.1 or 0.5 mM IPTG.

#### Protein purification

TniQ was overexpressed in BL21-AI cells. The cells were transformed with the pXT-H-TniQ expression plasmid. A single colony was used to inoculate the starter culture in LB broth and Carb. The following day, 15 mL of the overnight starter culture were used to inoculate 1.5 L of 2xYT medium containing Carb. Cells were incubated at 37 °C with shaking until the cell density reached OD_600_ 0.4. The temperature was then reduced to 18 °C and overexpression was induced by 0.5 mM IPTG and 0.2% arabinose. Cells were then grown overnight (18-20 hr) and harvested by centrifugation (5000g) for 15 min (4 °C) and the cell pellet was snap frozen with liquid nitrogen. Pellet was then thawed at RT for 30 min. Next, the pellet was resuspended in lysis buffer (20 mM Tris-HCl pH 8.0, 500 mM NaCl, 5% glycerol, 0.1% NP-40, 2 mM PMSF, EDTA-free complete protease inhibitor cocktail (Roche) and 5 mM BME). Cells were sonicated and centrifuged at 48,380g for 30 min at 4 °C, and the resulting supernatant was collected. The lysate was applied to a gravity column with pre-equilibrated Strep-Tactin Sepharose resin (IBA Life Sciences). The column was washed with 15 column volumes of 20 mM Tris-HCl pH 8.0, 500 mM NaCl, 5% glycerol and 5 mM BME. The protein was eluted with 20 mM Tris-HCl pH 8.0, 500 mM NaCl, 5% glycerol, 5 mM BME and 5 mM d-desthiobiotin. The eluted fractions were verified by running 4–20% SDS–PAGE gels.

#### Mate-out transposition assay

This assay monitors the movement of a mini-transposon (right and left ends flanking a Kan resistance gene) from a donor plasmid in BW27783 cells to a fertility (F) plasmid (carrying a Spec resistance gene). The F plasmid was engineered to contain: (1) the full-length putative target methylase gene recognized by the “*Ca.* Heimdallarchaeota” element (F-target), (2) the region upstream and including the native TSD (F-UP), (3) the region downstream and including the native TSD (F-DOWN), (4) the methylase gene with a mimicked insertion (F-target::Tn), (4) or a control *lacZα* fragment. Transposition proteins are expressed from inducible plasmid vectors in donor cells. After protein induction, the F plasmid is mated into recipient CW51 cells (Nal and Rif resistant) and the cell mix is plated with appropriate antibiotics. Transposition frequency is determined as the percentage of recipient cells with the mini-transposon marker from the total of recipient cells that received the F plasmid.

More specifically, donor cells were co-transformed with plasmids encoding the target selector (TniQ) and core transposase machinery (TnsAB and TnsC), then plated onto LB agar with glucose, Carb, and Cam. Transformants were washed up into M9 minimal media with appropriate combinations of maltose, 0.1 mM IPTG, 0/0.002/0.2% arabinose, Carb, and Cam to an OD_600_ of 0.3. After a 24 hr incubation at 30 or 16°C with aeration, induced cells were washed and resuspended in LB glucose, then incubated for 2 hr in 37°C. Cells were then mixed with recipient (1:10) and allowed to mate for 90 min at 37°C on a culture wheel turning at the lowest speed. The cell mix was serially diluted, and the correct dilutions were plated in LB agar containing appropriate antibiotics. Cells were simultaneously plated in Spec, Nal, Rif LB agar plates to calculate the total number of transconjugants, and in Spec, Nal, Rif, Kan LB agar plates to calculate the number of transconjugants that received the Kan resistance gene via transposition.

#### Mapping insertions

Transposition was confirmed by screening for TSDs. Isolates were obtained for colony PCR from the mate-out assay plates containing Kan. Amplicons were generated using a pair of primers that flank the methylase gene on the F plasmid. PCR products were checked for size, purified, and sent for Sanger sequencing.

Illumina sequencing was used to map insertions of mini-transposons in the F plasmids of transconjugants. Colonies were washed from the surface of mate-out assay plates containing Kan and resuspended in LB Kan Spec Nal. F plasmids were isolated using ZR BAC DNA Miniprep Kit and sent for paired end Illumina whole-genome sequencing through SeqCenter, LLC (seqcenter.com). Left and right ends were identified in the returned FASTA files and their coordinates were plotted to the length of the F plasmid. Analysis of NGS data was done using custom Python and shell scripts (data and code available at github.com/jordanthesier/Tn7_across_archaea).

#### Electrophoretic mobility shift assay

Roughly 200bp of substrate for the upstream and downstream methylase were generated by amplifying from a gblock of the reassembled “*Ca*. Heimdallarchaeota” methylase target gene. Amplicons were purified on a 2% agarose gel and eluted from gel slices.

Binding reactions were performed at 15 µL containing 20 mM Tris HCl pH 7.5, 100 mM NaCl, 10 µM ZnCl_2_, 6% glycerol, 5 mM BME, and 25ng of either the upstream or downstream substrate. Additionally, increasing concentrations of purified TniQ protein were added to each mixture followed by a 30 min incubation at 37 °C. After pre-running for 1 hr at 4°C, samples were run on a 5% non-denaturing polyacrylamide gel in 1× TBE buffer at 4°C. DNA was stained with SYBR^TM^ Gold and gel was imaged using a ChemiDocTM MP Imaging System (Bio-Rad).

#### Data Visualization

Similarity trees were visualized in iTOL (v7)^85^. Structural protein models were obtained from AlphaFold3^86^ and visualized in ChimeraX^87^, using Matchmaker for alignments.

#### Figs. S1 to S12

**Fig. S1.**
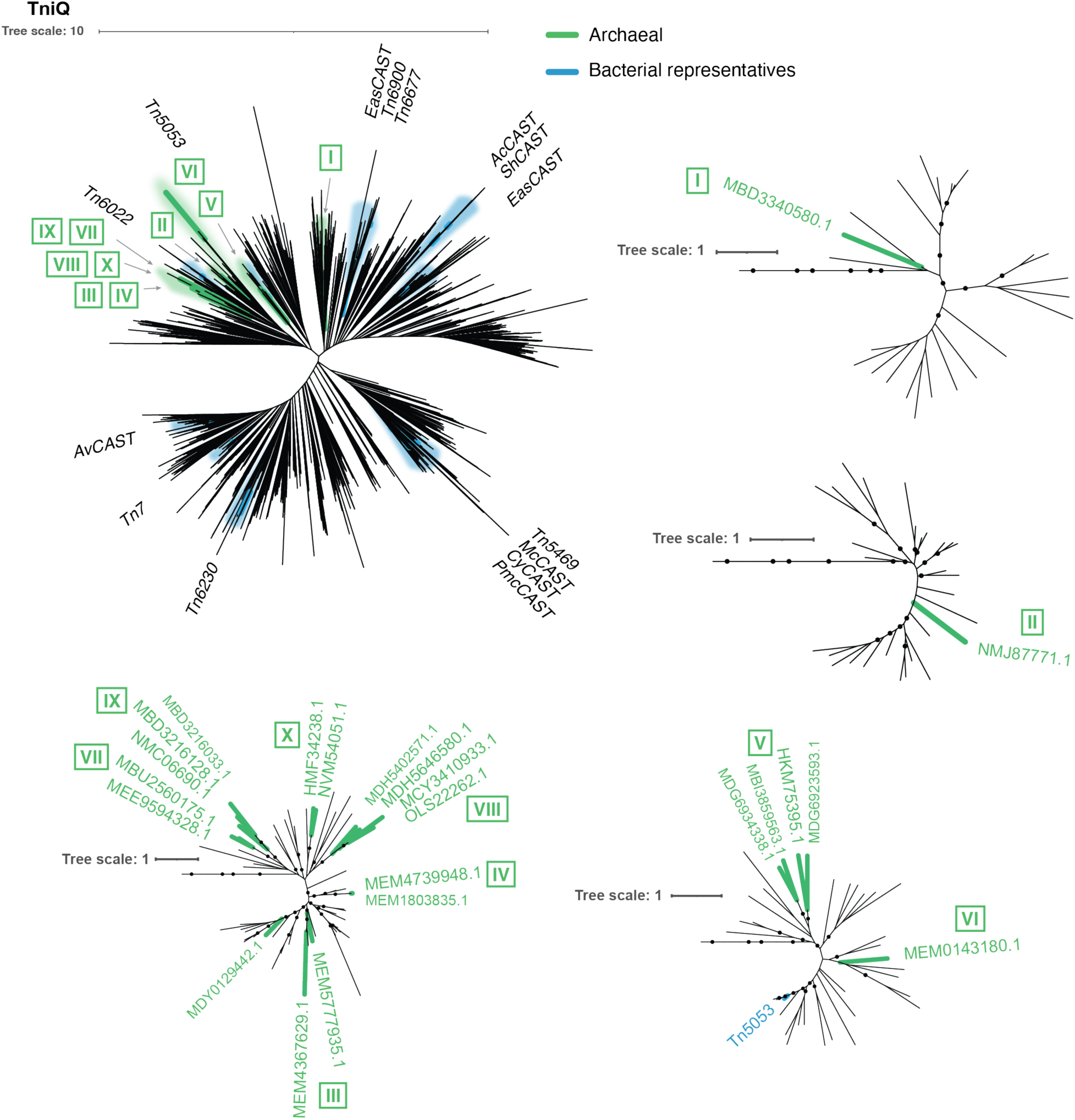
Distribution of archaeal TnsC proteins within an expanded Tn7 family dataset. A maximum likelihood tree of archaeal TnsC proteins supplemented with a comprehensive set of Tn7-like elements^42^. Green branches indicate archaeal proteins and blue branches indicate the 14 bacterial Tn7 family representatives. Roman numerals designate clade numbers. Clades with archaeal TnsC proteins are enlarged and proteins are labeled with NCBI accessions. Well-supported nodes (UFBoot ≥ 70%, SH-aLRT ≥ 0.8, aBayes ≥ 80%) are marked with black dots.

**Fig. S2.**
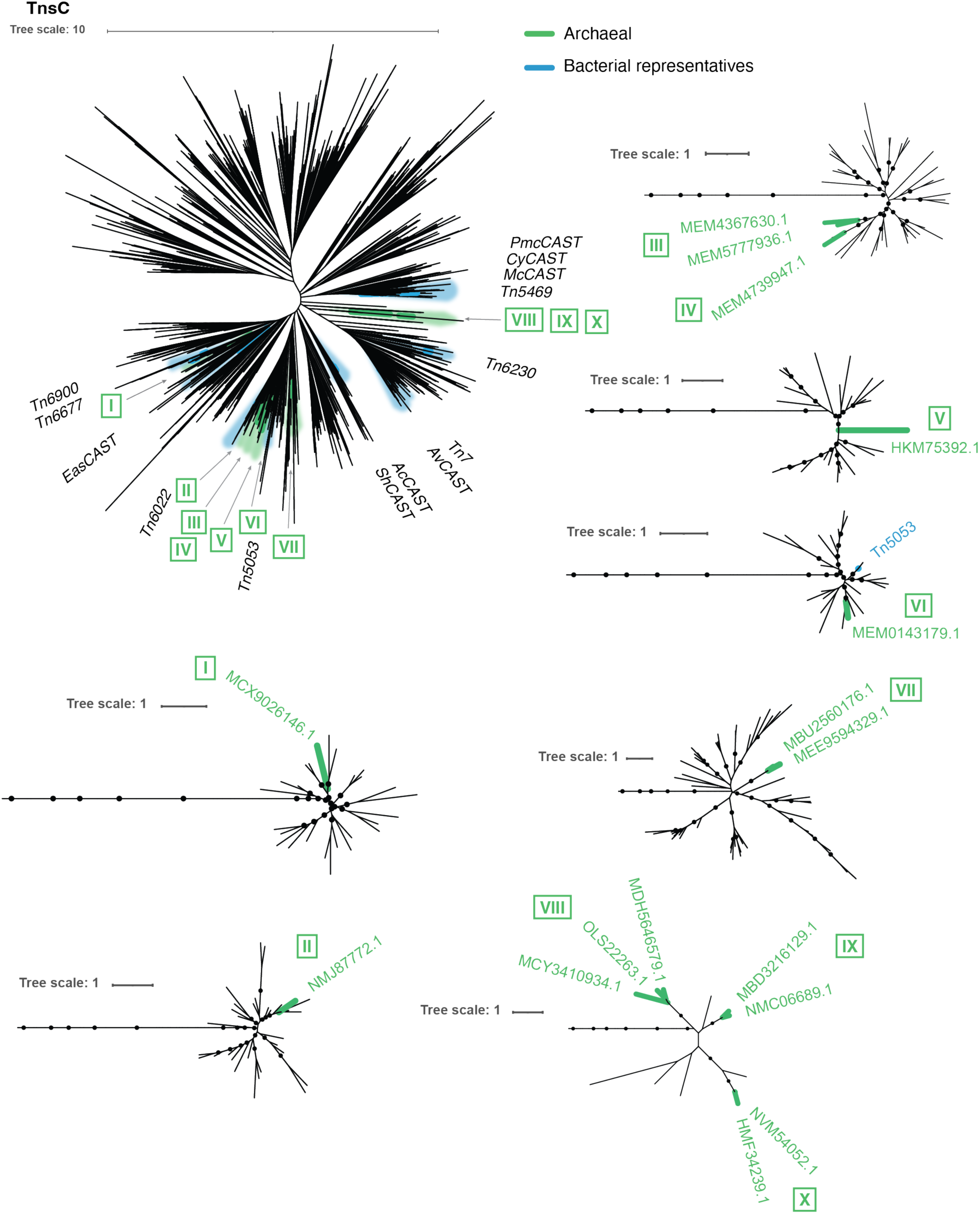
Distribution of archaeal TnsC proteins within an expanded Tn7 family dataset. A maximum likelihood tree of archaeal TniQ proteins supplemented with a comprehensive set of Tn7-like elements^42^. Green branches indicate archaeal proteins and blue branches indicate the 14 bacterial Tn7 family representatives. Roman numerals designate clade numbers. Clades with archaeal TniQ proteins are enlarged and proteins are labeled with NCBI accessions. Well-supported nodes (UFBoot ≥ 70%, SH-aLRT ≥ 0.8, aBayes ≥ 80%) are marked with black dots.

**Fig. S3.**
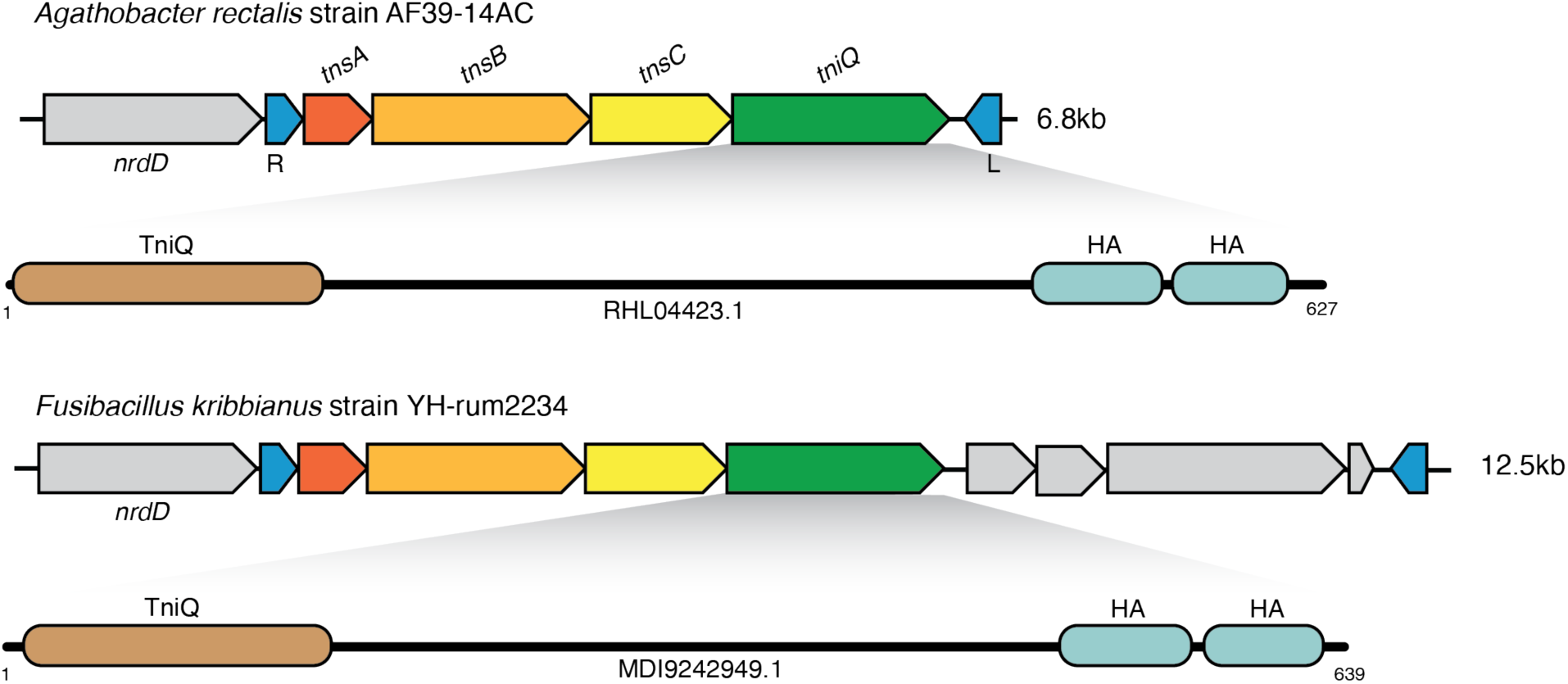
Bacterial *nrdD*-targeting Tn7-like elements. Gene configurations of two examples of Tn7-like transposons that target *nrdD* genes and domain architecture of their TniQ proteins. Transposition genes and transposons ends (L: left end; R: right end) are marked. Elements are labeled with the NCBI taxonomy of the host bacterium. Contig lengths are shown to the left. Protein domains were identified using InterProScan and are shown as colored ovals labeled with domain names. Domain positions are scaled to their coordinates within each full-length protein (black horizontal lines). NCBI protein accessions are indicated below each schematic. Abbreviations: HA, helicase-associated. Profiles: TniQ (PF06527); HA (PF03457).

**Fig. S4.**
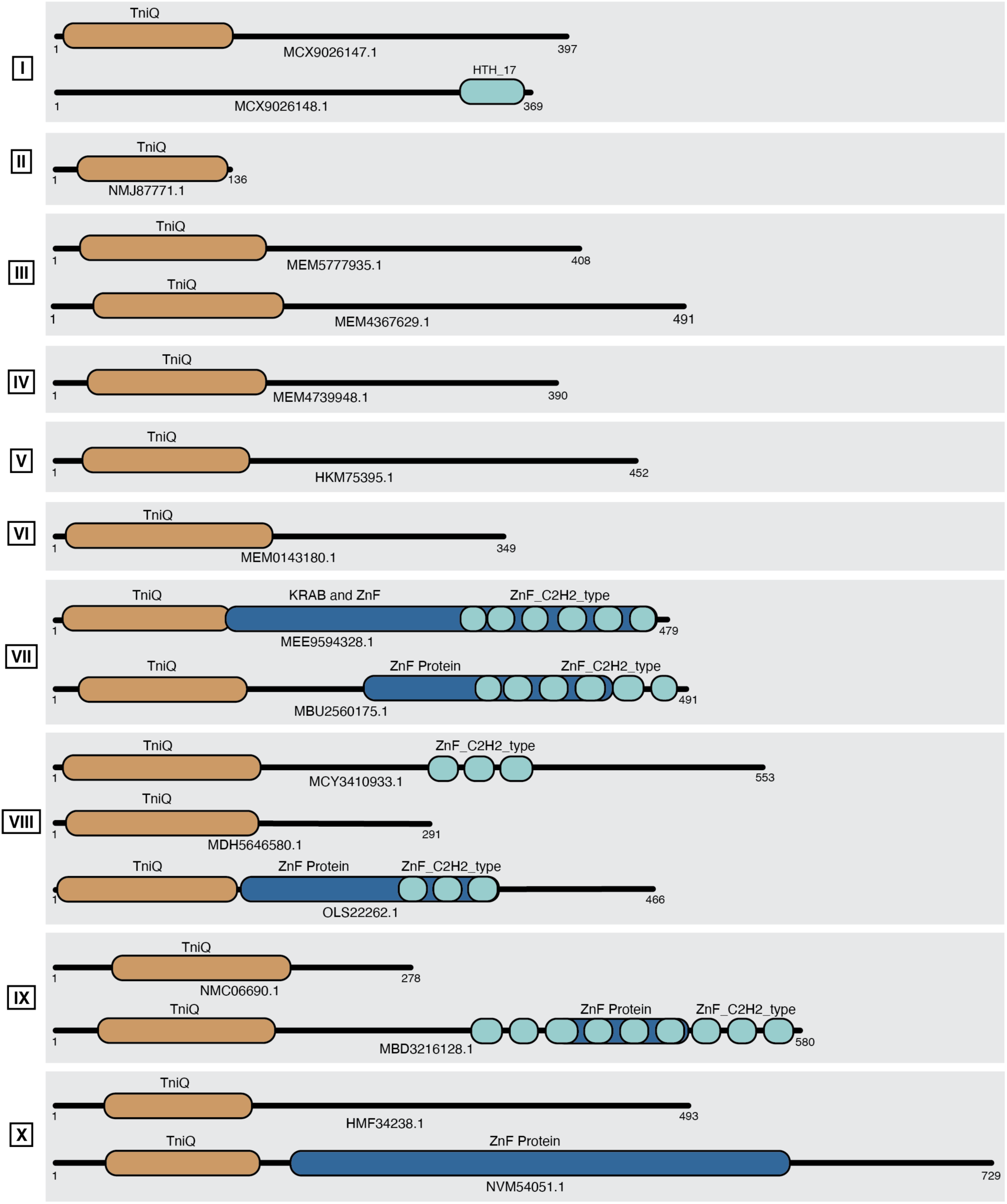
Domain architecture of archaeal TniQ proteins. Protein domains were identified using InterProScan and are shown as colored ovals labeled with domain names. Domain positions are scaled to their coordinates within each full-length protein (black horizontal lines). NCBI protein accessions are indicated below each schematic. Proteins are grouped based on archaeal clade, indicated by Roman numerals. Abbreviations: HTH, helix-turn-helix; ZnF, zinc finger; ZnF_C2H2_type, Cys_2_-His_2_ zinc finger type. Profiles: TniQ (PF06527); HTH_17 (PF12728), KRAB and ZnF (PTHR24379); ZnF Protein (PTHR24390, PTHR24404, PTHR24409); ZnF_C2H2_type (SM00355).

**Fig. S5.**
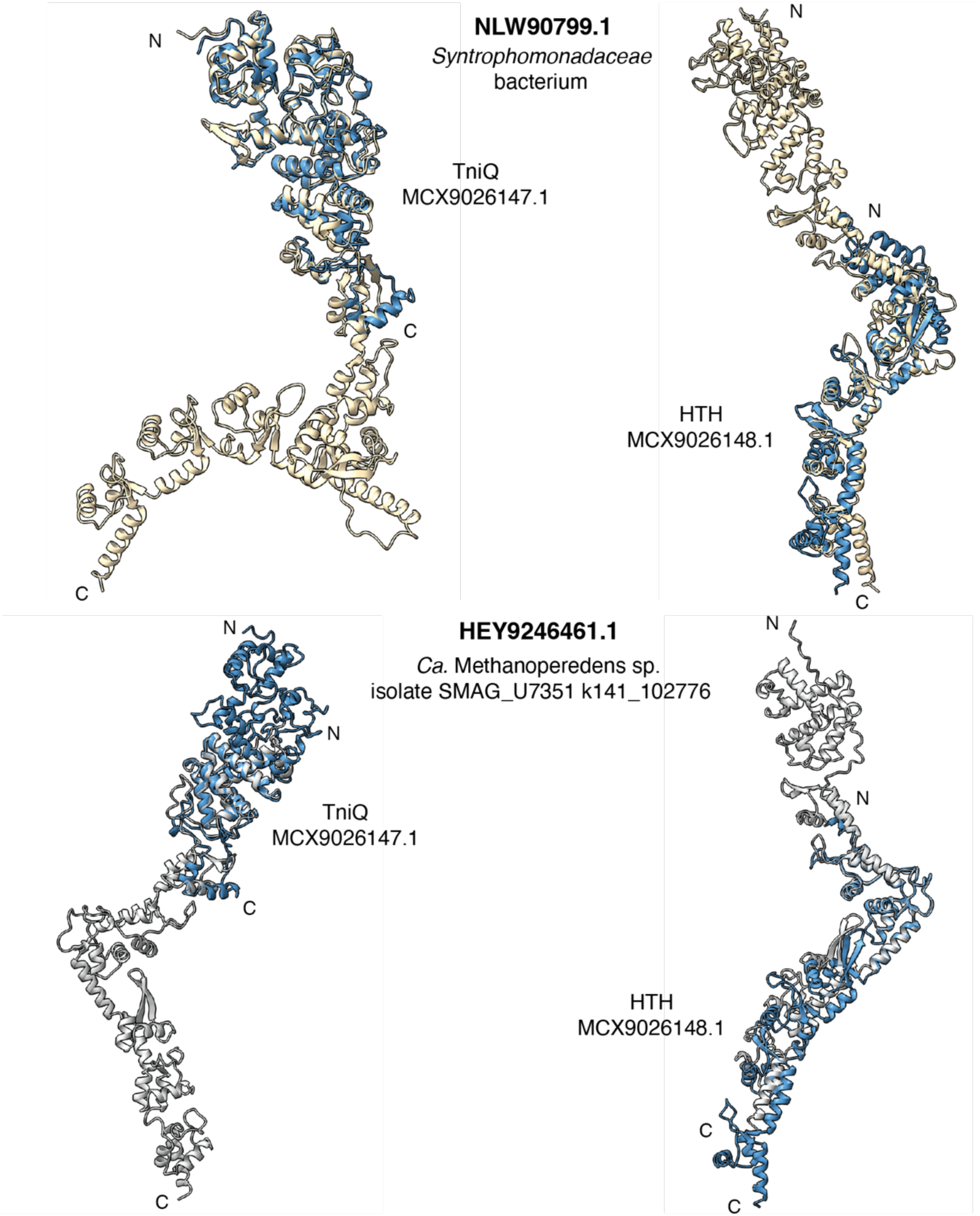
Clade I archaeal element homologs. AlphaFold3 models of the Clade I TniQ and helix-turn-helix (HTH) proteins aligned against their TniQ-HTH fusion homologs from bacteria (top) and archaea (bottom). TniQ alignments are shown on the left, and HTH alignments on the right. Clade I models are shown in blue; bacterial homolog is shown is beige; archaeal homolog is shown in grey. N- and C-termini are noted.

**Fig. S6.**
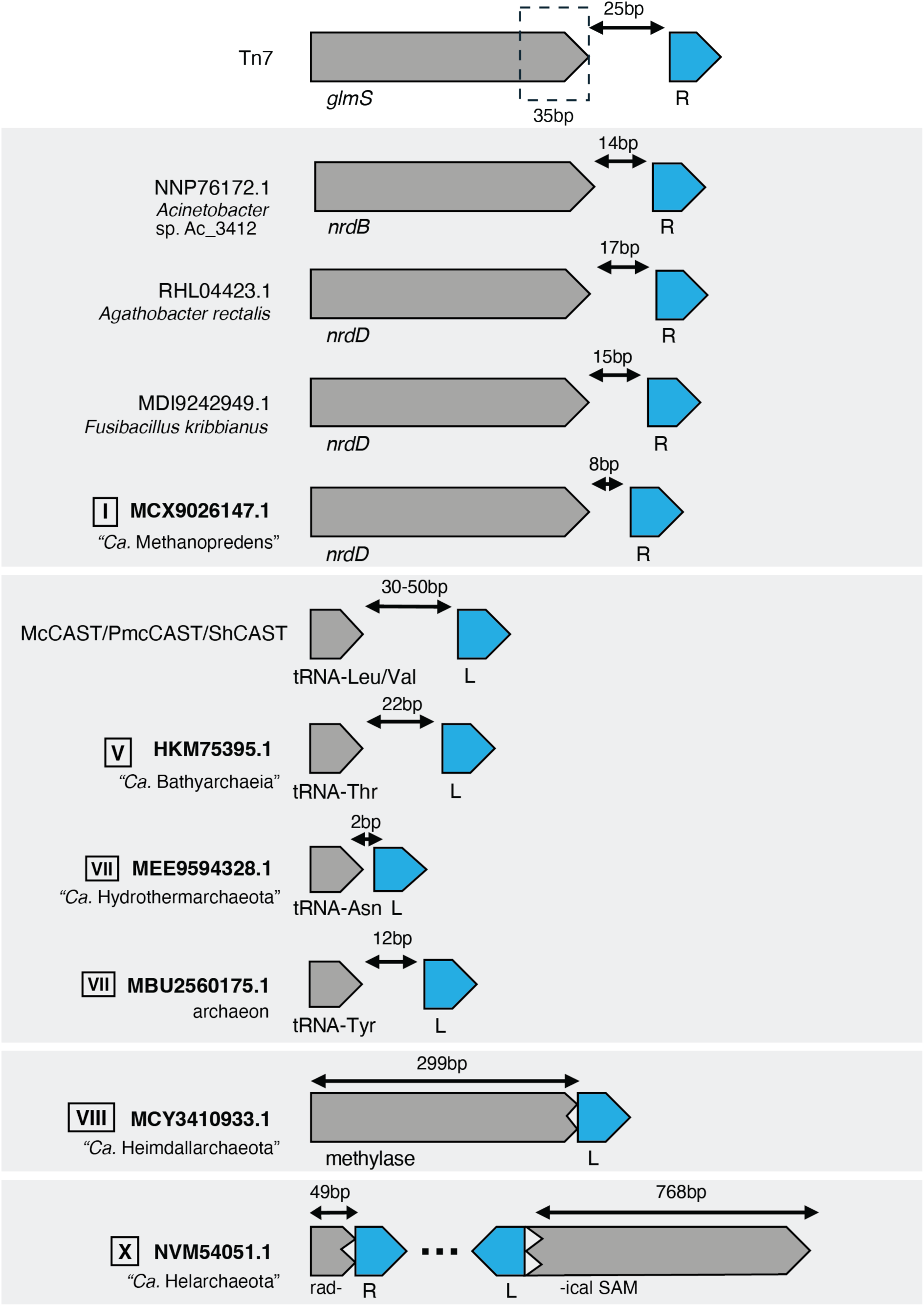
Spacing of Tn7 family transposon insertions. Gene configurations of archaeal and bacterial Tn7 family transposons are shown, highlighting the distance between transposon ends and adjacent target genes. Target genes are shown in gray and target-proximal transposon ends are shown in blue (L: left end; R: right end). The number of base pairs between the transposon end and the stop codon of the target gene is indicated. Elements are grouped by similar targets. Roman numerals represent clade numbers. TniQ NCBI accessions and host organisms are labeled. *Top*: Canonical Tn7 insertion is shown for reference; the dashed box indicates the TnsD/TniQ binding site.

**Fig. S7.**
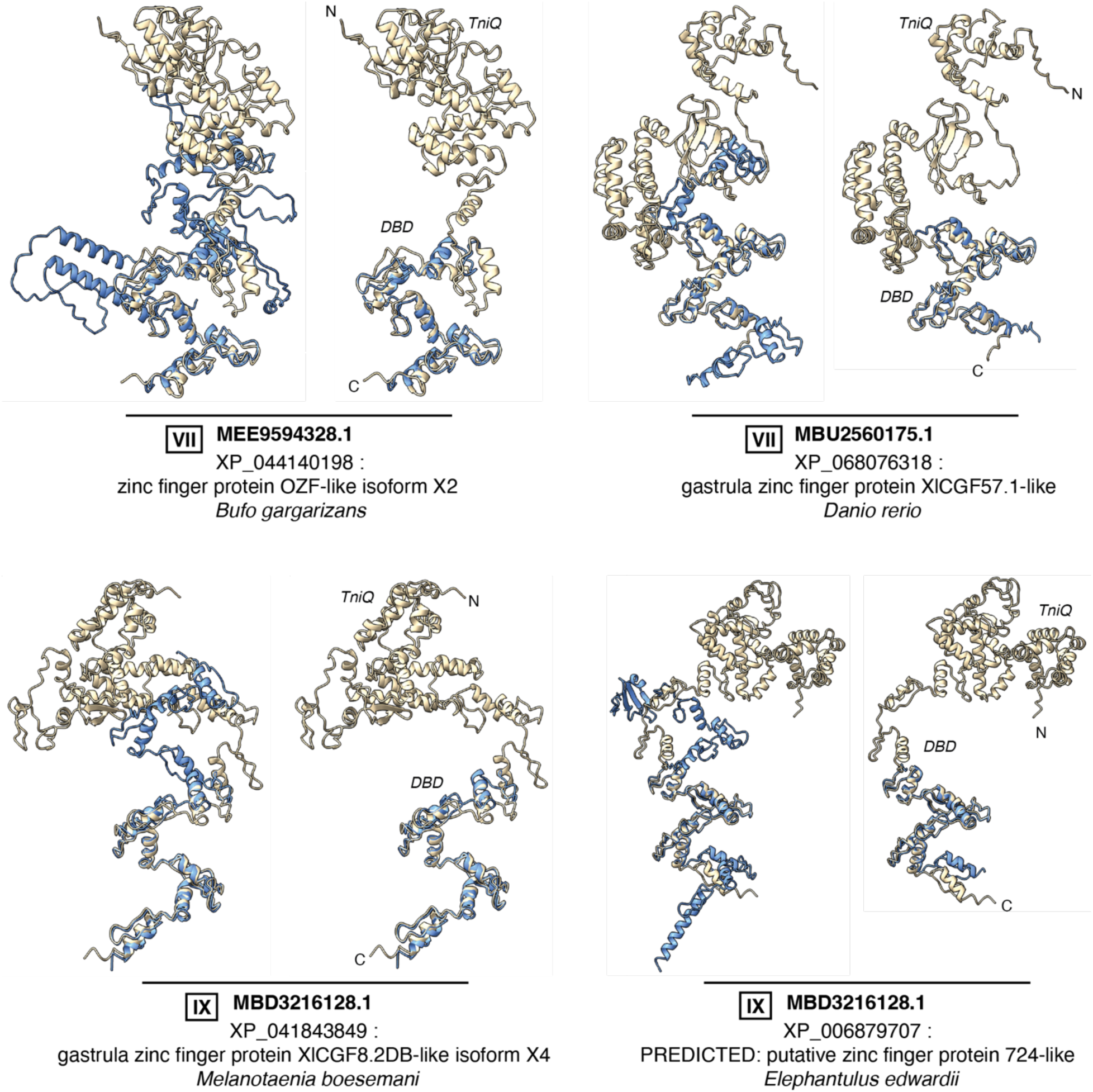
Predicted structural alignments of archaeal TniQ DNA-binding domains with eukaryotic homologs. AlphaFold3 models of archaeal TniQ proteins and their eukaryotic homologs found by BLAST. Each alignment includes NCBI protein accessions as well as a brief protein description and the host organism of the eukaryotic homolog. Roman numerals indicate archaeal clades. Archaeal TniQ models are shown in beige; eukaryotic homologs are shown in blue. For each alignment, the full-length eukaryotic homolog is displayed on the left, and a truncated version highlighting alignment with TniQ DNA-binding domain is shown on the right. N- and C-termini are noted on TniQs, as well as the TniQ domain and DNA-binding domain (DBD).

**Fig. S8.**
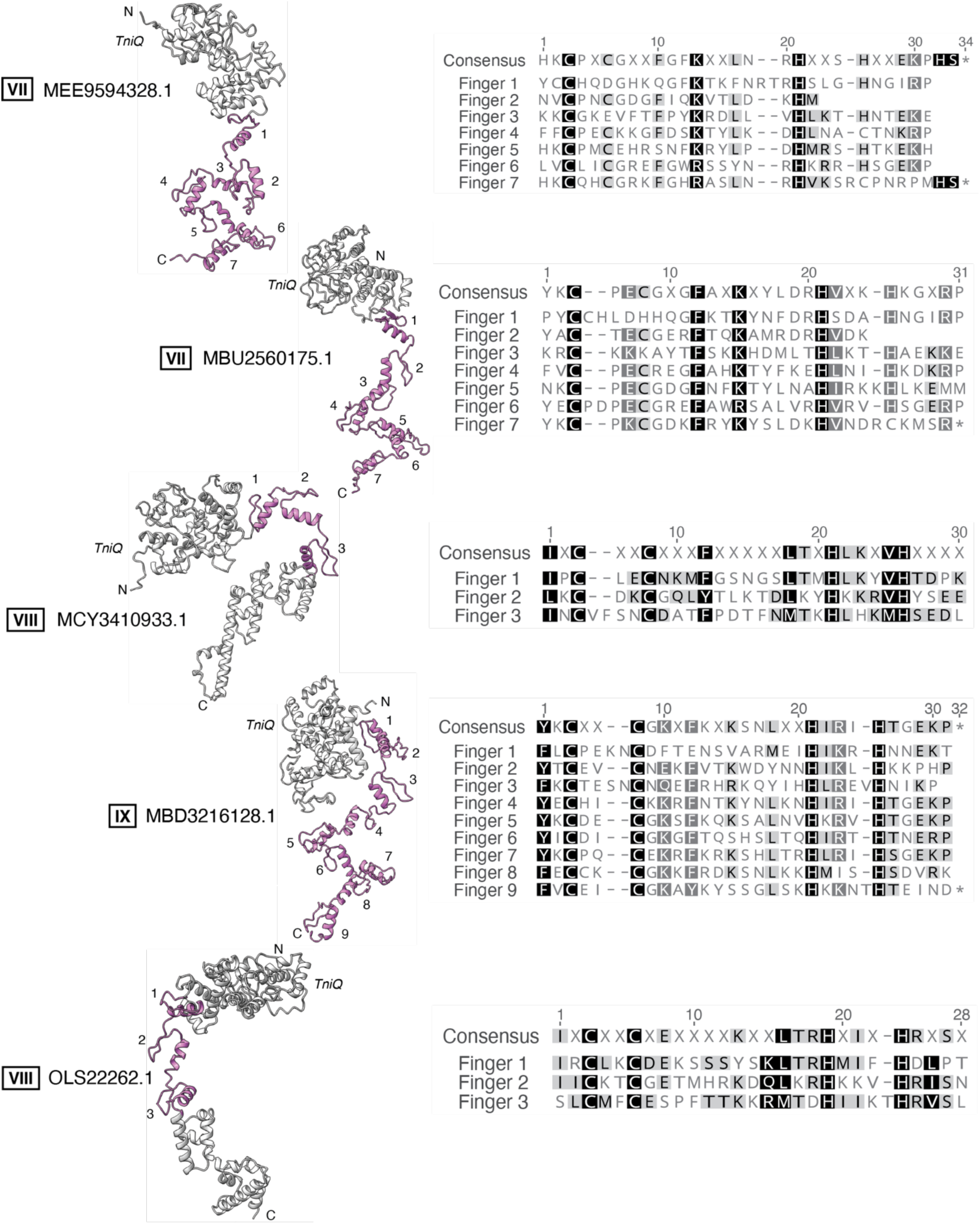
C2H2-ZnF motifs in archaeal TniQ proteins. *Left*: AlphaFold3 predicted structures of archaeal TniQ proteins. N- and C-termini are labeled, as well as TniQ domains. Cys_2_-His_2_ zinc fingers (C2H2-ZnFs) are shown in pink and numbered sequentially. NCBI TniQ accessions are noted with Roman numerals indicating archaeal clades. *Right*: Motif validation of each C2H2-ZnF. Finger numbers correspond with those shown on structural models. Darker backgrounds indicate higher similarity.

**Fig. S9.**
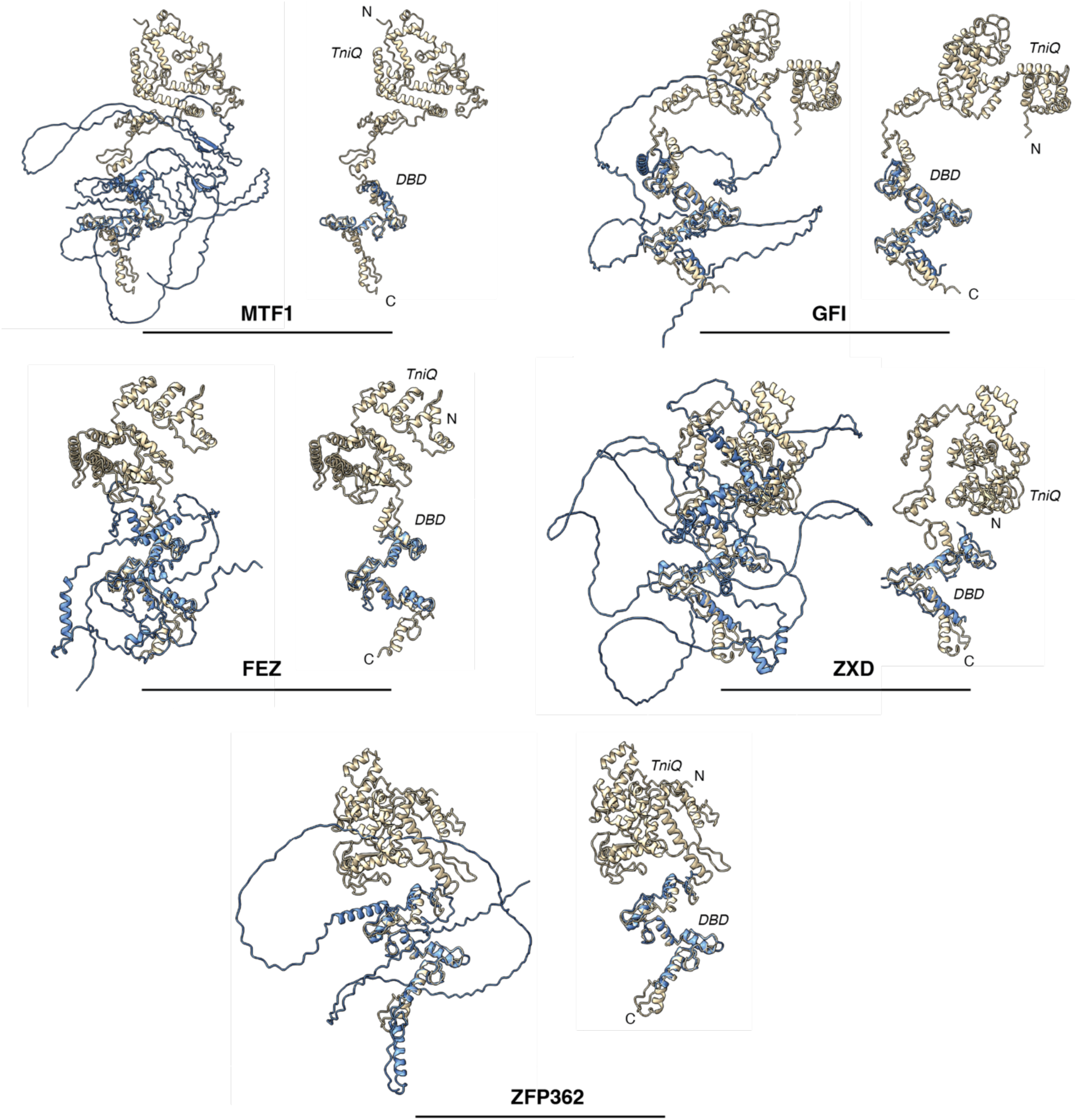
Predicted structural alignments of archaeal TniQ DNA-binding domain with human C2H2-ZnFs. AlphaFold3 models of the archaeal TniQ protein MBD3216128.1 and *Homo sapiens* representatives of five Cys_2_-His_2_ zinc finger (C2H2-ZnFs) proteins^71^. Archaeal TniQ model is shown in beige; *Homo sapiens* models are shown in blue. For each alignment, the full-length *Homo sapiens* C2H2-ZnF is displayed on the left, and a truncated version highlighting alignment with TniQ DNA-binding domain is shown on the right. The corresponding C2H2-ZnF family is noted below. N- and C-termini are noted on TniQs, as well as the TniQ domain and DNA-binding domain (DBD).

**Fig. S10.**
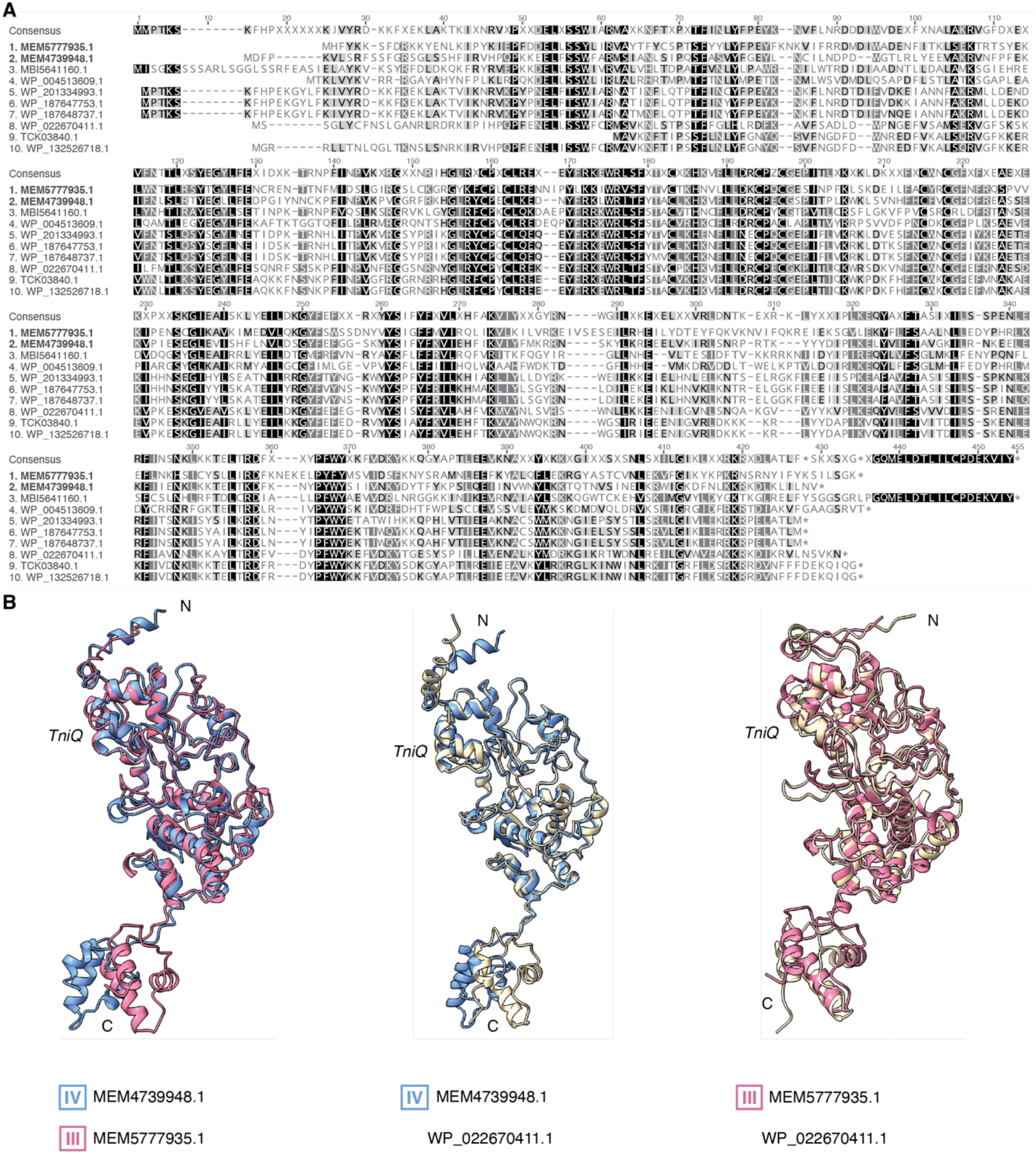
Clade III and IV TniQ bacterial homologs. **(A)** Alignment of protein sequences of MEM5777935.1 (Clade III), MEM4739948.1 (Clade IV), and eight bacterial TniQ homologs. Darker backgrounds indicate higher similarity. NCBI protein accessions are indicated to the left. **(B)** AlphaFold3 models and alignments of MEM5777935.1 (pink), MEM4739948.1 (blue), and a bacterial TniQ homolog WP_022670411.1 (beige). Roman numerals indicate archaeal clades.

**Fig. S11.**
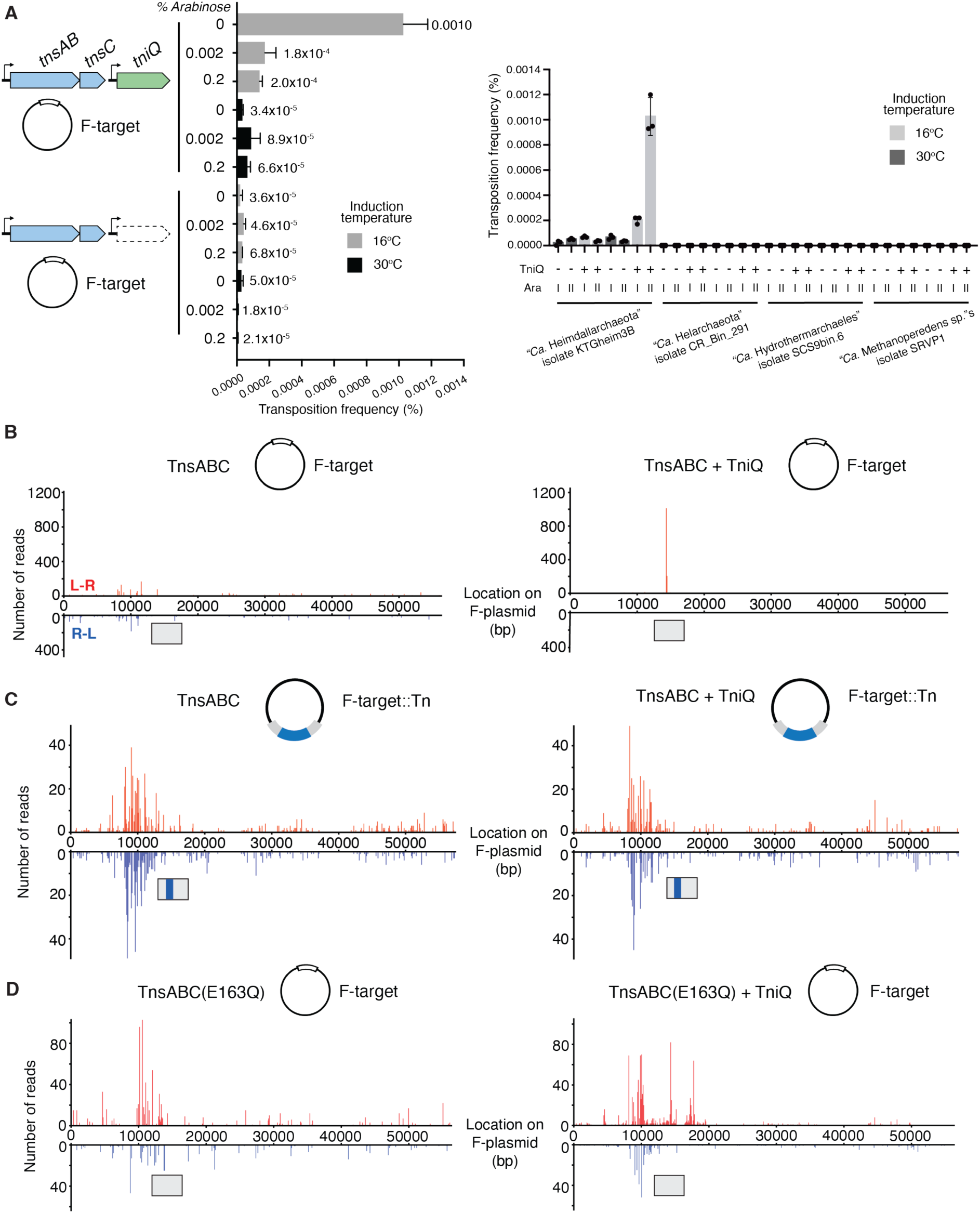
Expanded experimental conditions for archaeal transposition assays. **(A)** *Left*: Optimization of mate-out assay with the “*Ca*. Heimdallarchaeota” transposon. Proteins were induced with 0.1 mM IPTG and either 0, 0.002, or 0.2% arabinose at 30 or 16 °C. Data are shown as mean + SD, n = 3. *Right*: Transposition frequencies of three additional archaeal elements, with the “*Ca*. Heimdallarchaeota” element as a control, as measured by mate-out assay. Proteins were induced with 0.1 mM IPTG and either 0.2% (I) or 0% (II) arabinose (Ara) at 30 or 16 °C. Trials with and without TniQ expression are denoted with “+” and “-”, respectively. Data are shown as mean + SD, n = 3. **(B-D)** Insertion profiles across entire F plasmids from various mate-out assays with the “*Ca*. Heimdallarchaeota” element. Proteins expressed and F plasmid derivatives used are indicated above each graph. N- and C-termini are noted, as well as the TniQ domain.

**Fig. S12.**
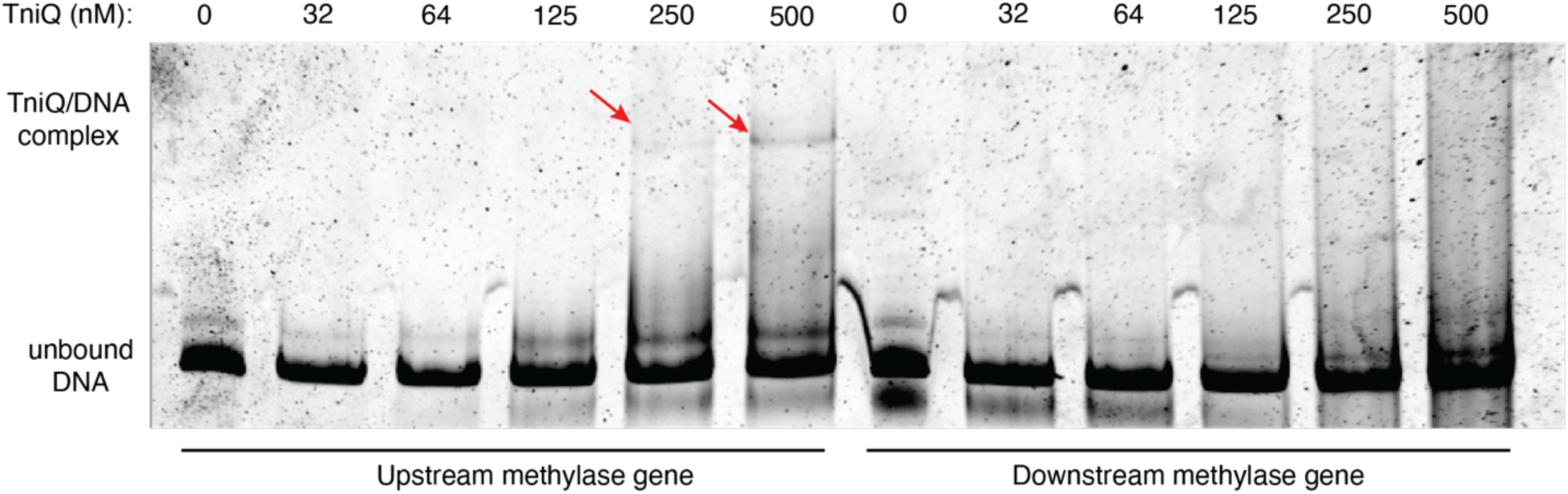
The archaeal TniQ recognizes the upstream fragment of the host methylase gene. Gel electrophoresis results of EMSA with increasing concentrations of purified Heimdall TniQ incubated with 25ng of either the upstream or downstream DNA fragment derived from the host methylase gene. The arrow indicates the position of the shifted DNA band predicted to be bound by TniQ.

#### Tables S1 to S8

**Table S1. NCBI BLAST output of MAGPurify marker genes and other archaeal-supporting markers on *tniQ*-containing contigs.** BLAST output of MAGPurify archaeal marker genes (top) and other archaeal-supporting genes (bottom) to support the archaeal origin of Tn7-like proteins. Includes identifier, accession, description, scientific name, query coverage, percent identity, e-value, and accession length.

**Table S2. Archaeal TniQ protein IDs and associated metadata.** All TniQ proteins found in this study and their associated metadata. Includes TniQ accessions, TnsC accessions, clade numbers, assembly id, contig id, NCBI organism name, GTDB taxonomy, CheckM2 results, and MAGPurify results.

**Table S3. NCBI BLAST output of protein sequences C-terminal to TniQ domains of archaeal TniQ proteins.** BLAST output of C-terminal protein sequences in TniQs to support examine DNA-binding domains. Includes identifier, accession, description, scientific name, query coverage, percent identity, e-value, and accession length.

**Table S4. NCBI BLAST output of *Homo sapiens* C2H2-ZnFs.** BLAST output of human representatives of five C2H2 families from Seetharam and Stuart, 2013 to support to confirm human origins. Includes identifier, accession, description, scientific name, query coverage, percent identity, e-value, and accession length.

**Table S5. NCBI Batch Conserved Domain Search output for targets and cargo of putative archaeal Tn7-like elements.** Batch Conserved Domain Search output for functional annotation of Tn7 target (top) and cargo (bottom) genes. Includes coordinates, e-values, bitscores, hit names, and descriptions.

**Table S6. EggNOG-mapper output of archaeal contigs with putative complete Tn7-like elements.** EggNOG-mapper output for functional annotation of genes on contigs with putative complete Tn7-like elements. Includes e-values, scores, maximum taxonomic annotation levels, COG categories, descriptions, KEGG pathways/reactions, and Pfam profiles.

**Table S7. Plasmids and strains used in this study.** Description of plasmids and strains including names, sources, markers, and sequences.

**Table S8. List of oligonucleotides used in this study.** Description of oligonucleotides including names and sequences.

## References

1 Hsieh, S. C. & Peters, J. E. Natural and Engineered Guide RNA-directed Transposition with CRISPR-Associated Tn7-like Transposons. Annu Rev Biochem (2024). 10.1146/annurev-biochem-030122-041908

2 Peters, J. E. Tn7. Microbiol Spectr 2, 1–20 (2014). 10.1128/microbiolspec.MDNA3-0010-2014

3 Sarnovsky, R., May, E. W. & Craig, N. L. The Tn7 transposase is a heteromeric complex in which DNA breakage and joining activities are distributed between di[erent gene products. EMBO J. 15, 6348–6361 (1996).

4 May, E. W. & Craig, N. L. Switching from cut-and-paste to replicative Tn7 transposition. Science 272, 401–404 (1996).

5 Lichtenstein, C. & Brenner, S. Site-specific properties of Tn7 transposition into the E. coli chromosome. Mol Gen Genet 183, 380–387 (1981). 10.1007/BF00270644

6 Arciszewska, L. K. & Craig, N. L. Interaction of the Tn7-encoded transposition protein TnsB with the ends of the transposon. Nucleic Acids Res 19, 5021–5029 (1991). 10.1093/nar/19.18.5021

7 Kaczmarska, Z. et al. Structural basis of transposon end recognition explains central features of Tn7 transposition systems. Mol Cell 82, 2618–2632.e2617 (2022). 10.1016/j.molcel.2022.05.005

8 Shen, Y. et al. Assembly of the Tn7 targeting complex by a regulated stepwise process. Mol Cell 84, 2368–2381 e2366 (2024). 10.1016/j.molcel.2024.05.012

9 Shen, Y. et al. Structural basis for DNA targeting by the Tn7 transposon. Nat Struct Mol Biol 29, 143–151 (2022). 10.1038/s41594-022-00724-8

10 Park, J. U., Tsai, A. W., Chen, T. H., Peters, J. E. & Kellogg, E. H. Mechanistic details of CRISPR-associated transposon recruitment and integration revealed by cryo-EM. Proc Natl Acad Sci U S A 119, e2202590119 (2022). 10.1073/pnas.2202590119

11 Choi, K. Y., Li, Y., Sarnovsky, R. & Craig, N. L. Direct interaction between the TnsA and TnsB subunits controls the heteromeric Tn7 transposase. Proceedings of the National Academy of Sciences 110, E2038–2045 (2013).

12 Park, J. et al. Structures of the holo CRISPR RNA-guided transposon integration complex. Nature 613, 775–782 (2022). doi.org/10.1038/s41586-022-05573-5

13 Baker, B. J. et al. Diversity, ecology and evolution of Archaea. Nat Microbiol 5, 887–900 (2020). 10.1038/s41564-020-0715-z

14 Eme, L. et al. Inference and reconstruction of the heimdallarchaeial ancestry of eukaryotes. Nature 618, 992–999 (2023). 10.1038/s41586-023-06186-2

15 Liu, Y. et al. Expanded diversity of Asgard archaea and their relationships with eukaryotes. Nature 593, 553–557 (2021). 10.1038/s41586-021-03494-3

16 Zhang, J. et al. Deep origin of eukaryotes outside Heimdallarchaeia within Asgardarchaeota. Nature (2025). 10.1038/s41586-025-08955-7

17 Wu, F. et al. Unique mobile elements and scalable gene flow at the prokaryote-eukaryote boundary revealed by circularized Asgard archaea genomes. Nat Microbiol 7, 200–212 (2022). 10.1038/s41564-021-01039-y

18 Parks, A. R. & Peters, J. E. Transposon Tn7 is widespread in diverse bacteria and forms genomic islands. Journal of Bacteriology 189, 2170–2173 (2007). 10.1128/JB.01536-06

19 Parks, A. R. & Peters, J. E. Tn7 elements: engendering diversity from chromosomes to episomes. Plasmid 61, 1–14 (2009).

20 Benler, S. et al. Cargo Genes of Tn7-Like Transposons Comprise an Enormous Diversity of Defense Systems, Mobile Genetic Elements, and Antibiotic Resistance Genes. mBio 12, e0293821 (2021). 10.1128/mBio.02938-21

21 Chaumeil, P. A., Mussig, A. J., Hugenholtz, P. & Parks, D. H. GTDB-Tk v2: memory friendly classification with the genome taxonomy database. Bioinformatics 38, 5315–5316 (2022). 10.1093/bioinformatics/btac672

22 Shaw, J. & Yu, Y. W. Fast and robust metagenomic sequence comparison through sparse chaining with skani. Nat Methods 20, 1661–1665 (2023). 10.1038/s41592-023-02018-3

23 Price, M. N., Dehal, P. S. & Arkin, A. P. FastTree 2--approximately maximum-likelihood trees for large alignments. PLoS One 5, e9490 (2010). 10.1371/journal.pone.0009490

24 Ondov, B. D. et al. Mash: fast genome and metagenome distance estimation using MinHash. Genome Biol 17, 132 (2016). 10.1186/s13059-016-0997-x

25 Matsen, F. A., Kodner, R. B. & Armbrust, E. V. pplacer: linear time maximum-likelihood and Bayesian phylogenetic placement of sequences onto a fixed reference tree. BMC Bioinformatics 11, 538 (2010). 10.1186/1471-2105-11-538

26 Jain, C., Rodriguez, R. L., Phillippy, A. M., Konstantinidis, K. T. & Aluru, S. High throughput ANI analysis of 90K prokaryotic genomes reveals clear species boundaries. Nat Commun 9, 5114 (2018). 10.1038/s41467-018-07641-9

27 Hyatt, D. et al. Prodigal: prokaryotic gene recognition and translation initiation site identification. BMC Bioinformatics 11, 119 (2010). 10.1186/1471-2105-11-119

28 Eddy, S. R. Accelerated Profile HMM Searches. PLoS Comput Biol 7, e1002195 (2011). 10.1371/journal.pcbi.1002195

29 Parks, D. H. et al. A complete domain-to-species taxonomy for Bacteria and Archaea. Nat Biotechnol 38, 1079–1086 (2020). 10.1038/s41587-020-0501-8

30 Parks, D. H. et al. GTDB: an ongoing census of bacterial and archaeal diversity through a phylogenetically consistent, rank normalized and complete genome-based taxonomy. Nucleic Acids Res 50, D785–D794 (2022). 10.1093/nar/gkab776

31 Parks, D. H. et al. A standardized bacterial taxonomy based on genome phylogeny substantially revises the tree of life. Nat Biotechnol 36, 996–1004 (2018). 10.1038/nbt.4229

32 Rinke, C. et al. A standardized archaeal taxonomy for the Genome Taxonomy Database. Nat Microbiol 6, 946–959 (2021). 10.1038/s41564-021-00918-8

33 Nayfach, S., Shi, Z. J., Seshadri, R., Pollard, K. S. & Kyrpides, N. C. New insights from uncultivated genomes of the global human gut microbiome. Nature 568, 505–510 (2019). 10.1038/s41586-019-1058-x

34 Wu, D., Jospin, G. & Eisen, J. A. Systematic identification of gene families for use as “markers” for phylogenetic and phylogeny-driven ecological studies of bacteria and archaea and their major subgroups. PLoS One 8, e77033 (2013). 10.1371/journal.pone.0077033

35 Chklovski, A., Parks, D. H., Woodcroft, B. J. & Tyson, G. W. CheckM2: a rapid, scalable and accurate tool for assessing microbial genome quality using machine learning. Nat Methods 20, 1203–1212 (2023). 10.1038/s41592-023-01940-w

36 Parks, D. H. et al. Recovery of nearly 8,000 metagenome-assembled genomes substantially expands the tree of life. Nat Microbiol 2, 1533–1542 (2017). 10.1038/s41564-017-0012-7

37 Schoelmerich, M. C. et al. A widespread group of large plasmids in methanotrophic Methanoperedens archaea. Nat Commun 13, 7085 (2022). 10.1038/s41467-022-34588-9

38 Peters, J. E., Makarova, K. S., Shmakov, S. & Koonin, E. V. Recruitment of CRISPR-Cas systems by Tn7-like transposons. Proceedings of the National Academy of Sciences 114, E7358 (2017). 10.1073/pnas.1709035114

39 McBride, T. M., Cameron, S. C., Fineran, P. C. & Fagerlund, R. D. The biology and type I/III hybrid nature of type I-D CRISPR-Cas systems. Biochem J 480, 471–488 (2023). 10.1042/BCJ20220073

40 Makarova, K. S. et al. An updated evolutionary classification of CRISPR-Cas systems. Nat Rev Microbiol 13, 722–736 (2015). 10.1038/nrmicro3569

41 Correa, A., 3rd et al. Novel mechanisms of diversity generation in Acinetobacter baumannii resistance islands driven by Tn7-like elements. Nucleic Acids Res 52, 3180–3198 (2024). 10.1093/nar/gkae129

42 Faure, G. et al. Modularity and diversity of target selectors in Tn7 transposons. Mol Cell 83, 2122–2136.e2110 (2023). 10.1016/j.molcel.2023.05.013

43 Ma, B. et al. A genomic catalogue of soil microbiomes boosts mining of biodiversity and genetic resources. Nat Commun 14, 7318 (2023). 10.1038/s41467-023-43000-z

44 Li, Z., He, Y., Zhang, H., Qian, H. & Wang, Y. Biotransformations of arsenic in marine sediments across marginal slope to hadal zone. J Hazard Mater 480, 135955 (2024). 10.1016/j.jhazmat.2024.135955

45 Hsieh, S. C. & Peters, J. E. Discovery and characterization of novel type I-D CRISPR-guided transposons identified among diverse Tn7-like elements in cyanobacteria. Nucleic Acids Res 51, 765–782 (2022). 10.1093/nar/gkac1216

46 Hsieh, S.-C. & Peters, J. E. Tn7-CRISPR-Cas12K elements manage pathway choice using truncated repeat-spacer units to target tRNA attachment sites. bioRxiv (2021). 10.1101/2021.02.06.429022

47 Faure, G. et al. CRISPR-Cas in mobile genetic elements: counter-defence and beyond. Nat Rev Microbiol 17, 513–525 (2019). 10.1038/s41579-019-0204-7

48 Saito, M. et al. Dual modes of CRISPR-associated transposon homing. Cell 184, 2441–2453 e2418 (2021). 10.1016/j.cell.2021.03.006

49 Hartman, H. & Fedorov, A. The origin of the eukaryotic cell: a genomic investigation. Proc Natl Acad Sci U S A 99, 1420–1425 (2002). 10.1073/pnas.032658599

50 Zaremba-Niedzwiedzka, K. et al. Asgard archaea illuminate the origin of eukaryotic cellular complexity. Nature 541, 353–358 (2017). 10.1038/nature21031

51 Zhao, R. & Biddle, J. F. Helarchaeota and co-occurring sulfate-reducing bacteria in subseafloor sediments from the Costa Rica Margin. ISME Commun 1, 25 (2021). 10.1038/s43705-021-00027-x

52 Klompe, S. E. et al. Evolutionary and mechanistic diversity of Type I-F CRISPR-associated transposons. Mol Cell 82, 616–628.e615 (2022). 10.1016/j.molcel.2021.12.021

53 Contreras-Moreira, B., Sancho, J. & Angarica, V. E. Comparison of DNA binding across protein superfamilies. Proteins 78, 52–62 (2010). 10.1002/prot.22525

54 Thomm, M. Archaeal transcription factors and their role in transcription initiation. FEMS Microbiol Rev 18, 159–171 (1996). 10.1111/j.1574-6976.1996.tb00234.x

55 Armache, K. J., Mitterweger, S., Meinhart, A. & Cramer, P. Structures of complete RNA polymerase II and its subcomplex, Rpb4/7. J Biol Chem 280, 7131–7134 (2005). 10.1074/jbc.M413038200

56 Eloranta, J. J., Kato, A., Teng, M. S. & Weinzierl, R. O. In vitro assembly of an archaeal D-L-N RNA polymerase subunit complex reveals a eukaryote-like structural arrangement. Nucleic Acids Res 26, 5562–5567 (1998). 10.1093/nar/26.24.5562

57 Aravind, L. & Koonin, E. V. DNA-binding proteins and evolution of transcription regulation in the archaea. Nucleic Acids Res 27, 4658–4670 (1999). 10.1093/nar/27.23.4658

58 Perez-Rueda, E. & Janga, S. C. Identification and genomic analysis of transcription factors in archaeal genomes exemplifies their functional architecture and evolutionary origin. Mol Biol Evol 27, 1449–1459 (2010). 10.1093/molbev/msq033

59 Bell, S. D. & Jackson, S. P. Mechanism and regulation of transcription in archaea. Curr Opin Microbiol 4, 208–213 (2001). 10.1016/s1369-5274(00)00190-9

60 Tadepally, H. D., Burger, G. & Aubry, M. Evolution of C2H2-zinc finger genes and subfamilies in mammals: species-specific duplication and loss of clusters, genes and e[ector domains. BMC Evol Biol 8, 176 (2008). 10.1186/1471-2148-8-176

61 Messina, D. N., Glasscock, J., Gish, W. & Lovett, M. An ORFeome-based analysis of human transcription factor genes and the construction of a microarray to interrogate their expression. Genome Res 14, 2041–2047 (2004). 10.1101/gr.2584104

62 Schuh, R. et al. A conserved family of nuclear proteins containing structural elements of the finger protein encoded by Kruppel, a Drosophila segmentation gene. Cell 47, 1025–1032 (1986). 10.1016/0092-8674(86)90817-2

63 Bouhouche, N., Syvanen, M. & Kado, C. I. The origin of prokaryotic C2H2 zinc finger regulators. Trends Microbiol 8, 77–81 (2000). 10.1016/s0966-842x(99)01679-0

64 Malgieri, G. et al. The prokaryotic Cys2His2 zinc-finger adopts a novel fold as revealed by the NMR structure of Agrobacterium tumefaciens Ros DNA-binding domain. Proc Natl Acad Sci U S A 104, 17341–17346 (2007). 10.1073/pnas.0706659104

65 Guilliere, F. et al. Solution structure of an archaeal DNA binding protein with an eukaryotic zinc finger fold. PLoS One 8, e52908 (2013). 10.1371/journal.pone.0052908

66 Bonchuk, A. N. & Georgiev, P. G. C2H2 proteins: Evolutionary aspects of domain architecture and diversification. Bioessays 46, e2400052 (2024). 10.1002/bies.202400052

67 Wolfe, S. A., Nekludova, L. & Pabo, C. O. DNA recognition by Cys2His2 zinc finger proteins. Annu Rev Biophys Biomol Struct 29, 183–212 (2000). 10.1146/annurev.biophys.29.1.183

68 Iuchi, S. Three classes of C2H2 zinc finger proteins. Cell Mol Life Sci 58, 625–635 (2001). 10.1007/PL00000885

69 Nowick, K., Hamilton, A. T., Zhang, H. & Stubbs, L. Rapid sequence and expression divergence suggest selection for novel function in primate-specific KRAB-ZNF genes. Mol Biol Evol 27, 2606–2617 (2010). 10.1093/molbev/msq157

70 Russo, J. et al. Sequences encoding C2H2 zinc fingers inhibit polyadenylation and mRNA export in human cells. Sci Rep 8, 16995 (2018). 10.1038/s41598-018-35138-4

71 Seetharam, A. & Stuart, G. W. A study on the distribution of 37 well conserved families of C2H2 zinc finger genes in eukaryotes. BMC genomics 14, 420 (2013). 10.1186/1471-2164-14-420

72 Mitra, R., McKenzie, G. J., Yi, L., Lee, C. A. & Craig, N. L. Characterization of the TnsD-attTn7 complex that promotes site-specific insertion of Tn7. Mobile DNA 1, 18 (2010).

73 Stellwagen, A. & Craig, N. L. Avoiding Self: Two Tn7-encoded proteins mediate target immunity in Tn7 transposition. EMBO J. 16, 6823–6834 (1997).

74 DeBoy, R. & Craig, N. L. Tn7 transposition as a probe of *cis* interactions between widely separated (190 Kilobases Apart) DNA sites in the *Escherichia coli* chromosome. J. Bacteriol. 178, 6184–6191 (1996).

75 Stellwagen, A. E. Analysis of gain of function mutants of an ATP-dependent regulator of Tn7 transposition. J. Mol. Biol. 305, 633–642 (2001).

76 Stellwagen, A. & Craig, N. L. Gain-of-function mutations in TnsC, an ATP-dependent transposition protein which activates the bacterial transposon Tn7. Genetics 145, 573–585 (1997).

77 Hsieh, S. C. et al. Telomeric transposons are pervasive in linear bacterial genomes. Science 387, eadp1973 (2025). 10.1126/science.adp1973

78 Klompe, S. E., Vo, P. L. H., Halpin-Healy, T. S. & Sternberg, S. H. Transposon-encoded CRISPR–Cas systems direct RNA-guided DNA integration. Nature 571, 219–225 (2019). 10.1038/s41586-019-1323-z

79 Zhang, J. et al. Deep origin of eukaryotes outside Heimdallarchaeia within Asgardarchaeota. Nature (2025). 10.1038/s41586-025-08955-7

80 Zhang, D., Burroughs, A. M., Vidal, N. D., Iyer, L. M. & Aravind, L. Transposons to toxins: the provenance, architecture and diversification of a widespread class of eukaryotic e[ectors. Nucleic Acids Res 44, 3513–3533 (2016). 10.1093/nar/gkw221

81 Minakhina, S., Kholodii, G., Mindlin, S., Yurieva, O. & Nikiforov, V. Tn5053 family transposons are *res* site hunters sensing plasmidal *res* sites occupied by cognate resolvases. Mol. Microbiol. 33, 1059–1068 (1999).

82 Bailey, T. L. & Elkan, C. Fitting a mixture model by expectation maximization to discover motifs in biopolymers. Proc Int Conf Intell Syst Mol Biol 2, 28–36 (1994).

83 Cantalapiedra, C. P., Hernandez-Plaza, A., Letunic, I., Bork, P. & Huerta-Cepas, J. eggNOG-mapper v2: Functional Annotation, Orthology Assignments, and Domain Prediction at the Metagenomic Scale. Mol Biol Evol 38, 5825–5829 (2021). 10.1093/molbev/msab293

84 Huerta-Cepas, J. et al. eggNOG 5.0: a hierarchical, functionally and phylogenetically annotated orthology resource based on 5090 organisms and 2502 viruses. Nucleic Acids Res 47, D309–D314 (2019). 10.1093/nar/gky1085

85 Letunic, I. & Bork, P. Interactive Tree of Life (iTOL) v6: recent updates to the phylogenetic tree display and annotation tool. Nucleic Acids Res 52, W78–W82 (2024). 10.1093/nar/gkae268

86 Abramson, J. et al. Accurate structure prediction of biomolecular interactions with AlphaFold 3. Nature 630, 493–500 (2024). 10.1038/s41586-024-07487-w

87 Meng, E. C. et al. UCSF ChimeraX: Tools for structure building and analysis. Protein Sci 32, e4792 (2023). 10.1002/pro.4792

